# Inflammaging in aged tissues drives remodeling of the CD8^+^ T cell compartment

**DOI:** 10.1101/2025.07.11.664388

**Authors:** Irina Shchukina, Carlos J. Rodriguez-Hernandez, Heather S. Ruiz, Maksim Kleverov, Rachel L. Mintz, Katsutaka Mineura, Subhadra C. Gunawardana, Sunnie Hsiung, Veronika Vachova, Jan Kossl, Denis A. Mogilenko, Christopher G. Huckstep, Barbora Vander Wielen, David W. Piston, Takeshi Egawa, Daniel Kreisel, Gwendalyn J. Randolph, Maxim N. Artyomov

## Abstract

Aging profoundly reshapes the immune cell landscape, with particularly strong effects on CD8^+^ T cells, including a marked decline in naïve cells and the emergence of age-associated GZMK^+^ CD8^+^ T cells (T_AA_ cells). Although T_AA_ cells make up a significant fraction of the aged CD8^+^ T cell compartment, the pathway underlying their development remains unknown. In this study, we demonstrate that T_AA_ cell development is cell-extrinsic and requires antigen exposure within aged non-lymphoid tissues. Using a novel TNF^Δ69AU/+^ mouse model, we show that systemic low-grade inflammation, characteristic of inflammaging, accelerates CD8^+^ T cell aging and promotes early accumulation of T_AA_ cells. Through detailed analysis of T_AA_ cell heterogeneity, we identified a progenitor subpopulation enriched in the aged adipose tissue. Using heterochronic transplantation, we show that adipose tissue acts as a functional niche, supporting progenitor maintenance and driving the conversion of young CD8^+^ T cells into the aged phenotype. Taken together, our findings reveal how aging of non-lymphoid tissues orchestrates the reorganization of the CD8^+^ T cell compartment and highlight adipose tissue as a promising target for therapeutic strategies aimed at modulating immune aging.

## INTRODUCTION

Aging has a profound impact on the CD8⁺ T cell compartment in both mice and humans^1,2^. Hallmark features of immune aging include thymic involution^3^, a progressive decline in naïve T cells, and a concomitant accumulation of memory T cell subsets^4^. In our previous work using aged mice, we identified a novel population of CD8⁺ T cells that accumulates across multiple tissues with age^5^. These age-associated CD8⁺ T cells (T_AA_ cells) are distinct from conventional effector and memory subsets and are also increased in the peripheral blood of older humans. At the transcriptional level, T_AA_ cells are marked by high expression of *Gzmk*, which encodes granzyme K (GZMK) – a granzyme implicated in both cytolytic and non-cytolytic functions^6^, including promotion of pro-inflammatory responses^7,8^. T_AA_ cells also exhibit co-expression of activation and exhaustion signatures, including the canonical markers *Pdcd1* and *Tox*, detectable at both the RNA and protein levels. Accordingly, T_AA_ cells were initially identified by flow cytometry as PD1⁺TOX⁺ CD8⁺ T cells. Their accumulation is driven by the aged environment, indicating a cell-extrinsic origin of the phenotype. T_AA_ cells are highly clonal, and cells isolated from different tissues often share T cell receptor (TCR) sequences, suggesting that they recirculate between organs. However, the mechanisms underlying T_AA_ cell development remain largely unknown.

The interplay between organismal aging and the immune system has been investigated from multiple perspectives, with both population-level and functional alterations in immune cells well documented^9,10^. More recently, several studies have provided compelling evidence that immune dysfunction – particularly within the T cell compartment – can actively drive age-related pathologies. For example, a hematopoietic-specific defect in DNA repair was shown to elevate systemic inflammatory cytokines and induce the accumulation of senescence markers across multiple tissues^11^. In another model, T cell-specific disruption of mitochondrial function resulted in severe systemic deterioration, including cardiovascular, metabolic, and cognitive decline, ultimately leading to a markedly shortened lifespan^12^. More recently, targeted deletion of *Rip1* in T cells induced excessive apoptosis and drove widespread T cell depletion, thus modeling the age-associated loss of naïve T cells^13^. These mice also developed hallmarks of accelerated aging such as osteoporosis, tissue senescence, and reduced fitness.

These studies primarily employed genetic models with immune- or T cell-intrinsic defects that led to phenotypes resembling natural aging of the immune system, and subsequently assessed how these alterations affected systemic physiology, survival, and overall fitness. Collectively, they established that dysfunctional immune cells are sufficient to drive broad, physiologically detrimental outcomes. However, our previous work demonstrated that the reverse is also true: the aged environment shapes the composition and phenotype of the CD8⁺ T cells^5^. Specifically, we showed that young, phenotypically normal CD8⁺ T cells acquire age-associated characteristics after residing in an aged host for several weeks. These findings point to a feedback loop in which the aged environment actively drives remodeling of the T cell pool. Therefore, we sought to define the steps required for this conversion and identify the environmental factors responsible for promoting the aged CD8⁺ T cell phenotype.

In this study, we report that chronic low-grade inflammation is sufficient to accelerate the aging of the CD8⁺ T cell compartment. We show that both the development and frequency of T_AA_ cells are governed by the aged environment. Our findings demonstrate that exposure to an aged non-lymphoid tissue microenvironment is a necessary step in T_AA_ cell generation, and notably, that aged adipose tissue alone is sufficient to drive this process. We identify aged fat as a critical niche that supports the emergence and expansion of T_AA_ progenitor cells, highlighting a direct link between tissue-specific aging and immune remodeling.

## RESULTS

### T_AA_ cells differentiation requires exposure to an antigen

We previously demonstrated that exposure to an aged environment is sufficient to drive the development of T_AA_ cells, indicating that this phenotype does not arise from cell-intrinsic changes within CD8^+^ T cells from old mice^5^. In this model of *de novo* T_AA_ cell generation, CD8^+^ T cells were isolated from the spleens of young congenically labeled donors and transferred into aged recipients (Fig. 1a, Supplementary Fig. 1a). Two to four weeks post-transfer, a substantial subset of young donor cells – initially characterized by a PD1⁻TOX⁻ phenotype – acquired an aged PD1⁺TOX⁺ phenotype (Fig. 1b). Furthermore, the transcriptional profile and homing pattern of these converted donor cells closely mirrored those of the host-derived native T_AA_ cells^5^. This high degree of similarity confirms that the transfer model faithfully recapitulates the natural development of T_AA_ cells with aging, providing a powerful system for mechanistic studies.

**Fig. 1:**
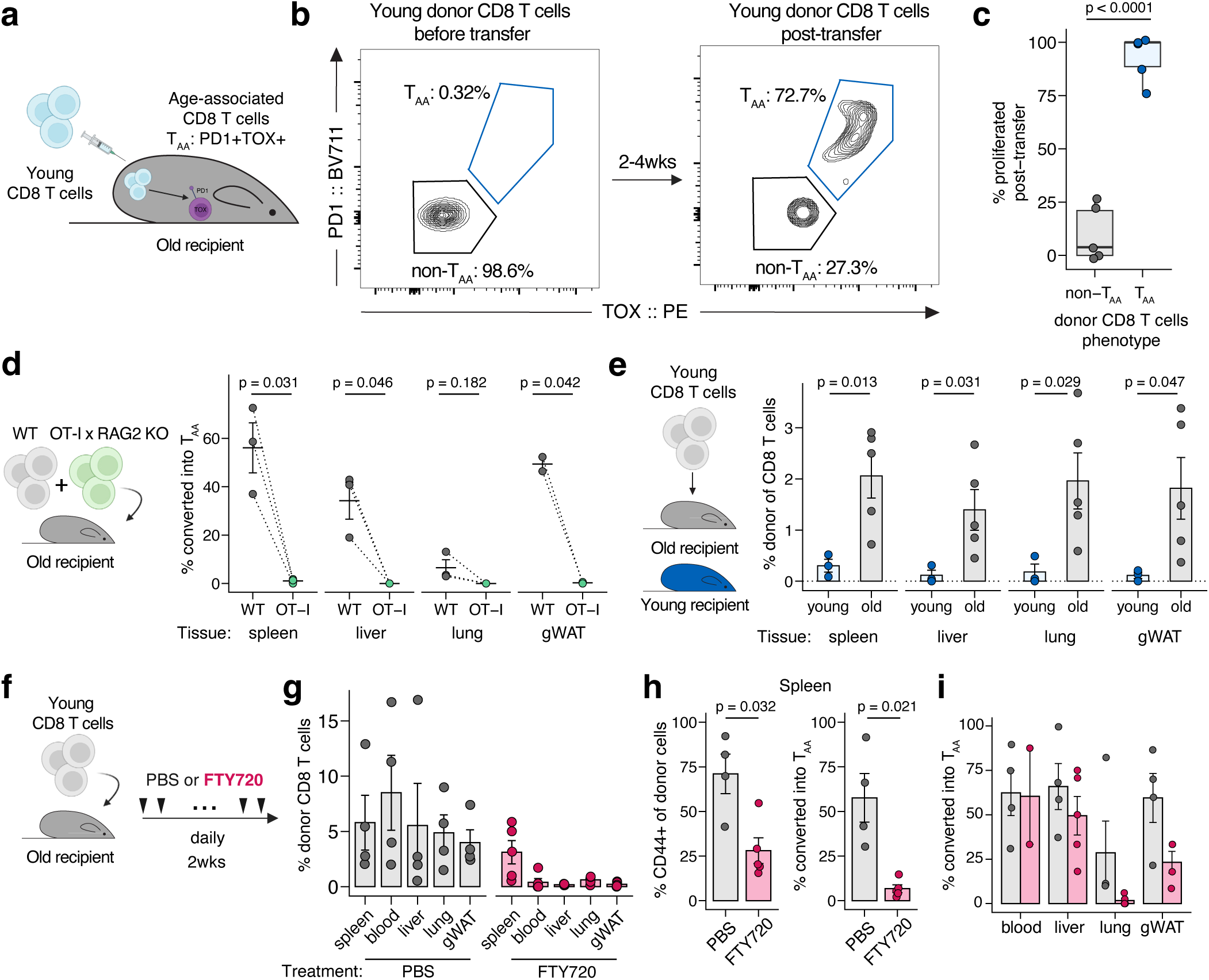
T_AA_ cells develop in aged peripheral tissues in an antigen-dependent manner. **a.** Transfer experiment design. **b.** PD1 and TOX expression in young donor CD8 T cells. Left: before transfer, right: spleen one month post-transfer. **c.** Frequency of donor cells in the blood that lost the Cell-Trace label 16 days post transfer. **d.** Frequency of WT or OT1 donor CD8 T cells that converted into T_AA_ cells one month post-transfer into old recipient. **e.** Infiltration of donor cells into tissues one month post-transfer into old or young recipients. **f.** Experiment design. **g.** Infiltration of donor cells into tissues two weeks post-transfer. **h.** Frequency of CD44^+^ donor cells (left) and PD1^+^TOX^+^ donor cells (right) in spleen. **i.** Frequency of leaked donor CD8 T cells converted into T_AA_ cells in tissues. Unpaired two-sample Student’s t-test **(c, e, h)**; paired two-sample Student’s t-test **(d)**. Data are mean ± s.e.m. for (**d, e, g-i**). Exact group sizes are provided in methods.

Given our prior observation that T_AA_ cells are highly clonal^5^, we hypothesized that young cells should undergo a proliferative stage during differentiation into T_AA_ cells. To test this, we stained young cells with CellTrace FarRed and transferred them into old hosts. Sixteen days later, we profiled the blood and found that those young cells that acquired the T_AA_ phenotype predominantly lost CellTrace (Fig. 1c), indicating that they had undergone several rounds of division. Notably, donor cells that retained the PD1^−^TOX^−^ phenotype showed minimal proliferation (Fig. 1c).

CD8^+^ T cells typically proliferate following activation through antigen recognition by the TCR^14^. To determine whether antigen exposure is required for T_AA_ cells differentiation, we utilized a TCR transgenic mouse model. We used OT-1 mice on a RAG2 knockout background to eliminate secondary TCR rearrangements, which naturally occur in approximately 15% of murine CD8^+^ T cells^15,16^. This approach ensured that all T cells remained specific to chicken ovalbumin – an antigen absent in naïve, unimmunized mice. We mixed OT-1 cells with polyclonal wild-type (WT) CD8^+^ T cells and transferred the mixture into old hosts. The difference between the genotypes was dramatic, with OT-1/RAG2KO cells failing to generate T_AA_ cells, in contrast to their WT counterparts exposed to the same environment (Fig. 1d). Thus, we established that T_AA_ cells are true antigen-experienced cells.

### T_AA_ cell differentiation requires access to old peripheral tissue

Next, we tested whether antigen recognition in an aged environment was sufficient to generate T_AA_ cells. We transferred OT-1 cells into young or old recipients and systemically delivered the SIINFEKL peptide recognized by these cells. While this approach successfully induced OT-1 cell activation, expansion, and memory formation, one-month post-transfer, the cells no longer expressed PD1 or TOX and failed to acquire a T_AA_ phenotype (Supplementary Fig. 1b). A similar result was observed when the immunization included an adjuvant (Supplementary Fig. 1c). This result suggested that CD8^+^ T cell activation and expansion following antigen recognition within an aged environment does not universally lead to the development of T_AA_ cells. To further investigate this possibility, we evaluated whether antigen-specific T_AA_ cells form following viral infection. Using the LCMV Armstrong model of acute infection, we found that antigen-specific T_AA_ cells did not develop in this context either (Supplementary Fig. 1d). Together, these results indicate that T_AA_ cell development depends on the context in which the antigen is presented.

Following immunization or acute infection, CD8^+^ T cells are typically primed in secondary lymphoid organs (SLOs)^17^. Given that in these models no T_AA_ cells were generated, we hypothesized that T_AA_ cell development might instead be driven by an atypical mode of priming, such as in peripheral tissues. Indeed, prior studies in young mice have shown that priming in the liver – a peripheral tissue allowing antigen recognition by naïve CD8⁺ T cells – can induce a dysfunctional transcriptional program^18^. Among peripheral sites, the liver appears to be uniquely permissive for CD8⁺ T cell priming in young mice^18^. In old mice, however, peripheral tissue access is altered globally. In our transfer experiments, we found that multiple old tissues were significantly more permissive to donor cell trafficking than young tissues, as evidenced by a higher number of donor cells detected in the periphery of old mice one month after transfer (Fig. 1e). Previously reported abnormal trafficking of CD8⁺ T cells in aging has been linked to age-related upregulation of tissue CXCL13^19^. However, we find that neither this chemokine nor the integrin CD49d, which is up-regulated on T_AA_ cells^5^, was required for T_AA_ cells development (Supplementary Fig. 1e-j).

To formally test whether tissue homing is required for the T_AA_ cells development, we transferred young CD8⁺ T cells into old recipients and treated half of the mice with FTY720 (Fig. 1f), a drug that prevents T cell egress from SLOs by blocking their re-entry into circulation^20^. As expected, FTY720 treatment resulted in a near-complete absence of donor cells in peripheral tissues (Fig. 1g). Surprisingly, the donor cells retained in the spleens of old mice did not acquire the T_AA_ phenotype (Fig. 1h), suggesting that differentiation into T_AA_ cells occurs outside of SLOs, within non-lymphoid tissues. Additionally, these spleen-retained cells largely maintained a CD44⁻ naïve phenotype (Fig. 1h), indicating that both antigen exposure and the necessary environmental cues occur in the periphery. Notably, the small number of donor cells that escaped into peripheral tissues in FTY720-treated mice did acquire the T_AA_ phenotype (Fig. 1i). Taken together, these data establish that access to peripheral tissues and priming outside of SLOs are required for T_AA_ cell development, highlighting the critical role of the aged tissue microenvironment in shaping CD8⁺ T cell phenotypes.

### T_AA_ cell abundance reflects variation in CD8⁺ T cell aging across individual mice

With accumulating data on T_AA_ cell abundance across many mice, we noticed that their frequencies in different tissues of the same mouse appeared to be correlated. For example, mice with a higher percentage of T_AA_ cells in the spleen also tended to have elevated levels in the liver. This pattern suggested the existence of organism-wide determinants influencing CD8⁺ T cell aging in individual mice. To investigate this more systematically, we analyzed a larger cohort of aged mice (Fig. 2a). We observed substantial inter-individual variability in the abundance of splenic T_AA_ cells (Fig. 2b). On average, approximately 34% of CD8⁺ T cells in the aged spleen exhibited the T_AA_ phenotype, but this proportion ranged widely – from as low as 11% to nearly 80%. As hypothesized, mice with high frequencies of splenic T_AA_ cells also showed similarly elevated levels in other organs (Fig. 2c), supporting the idea of systemic regulation.

**Fig. 2:**
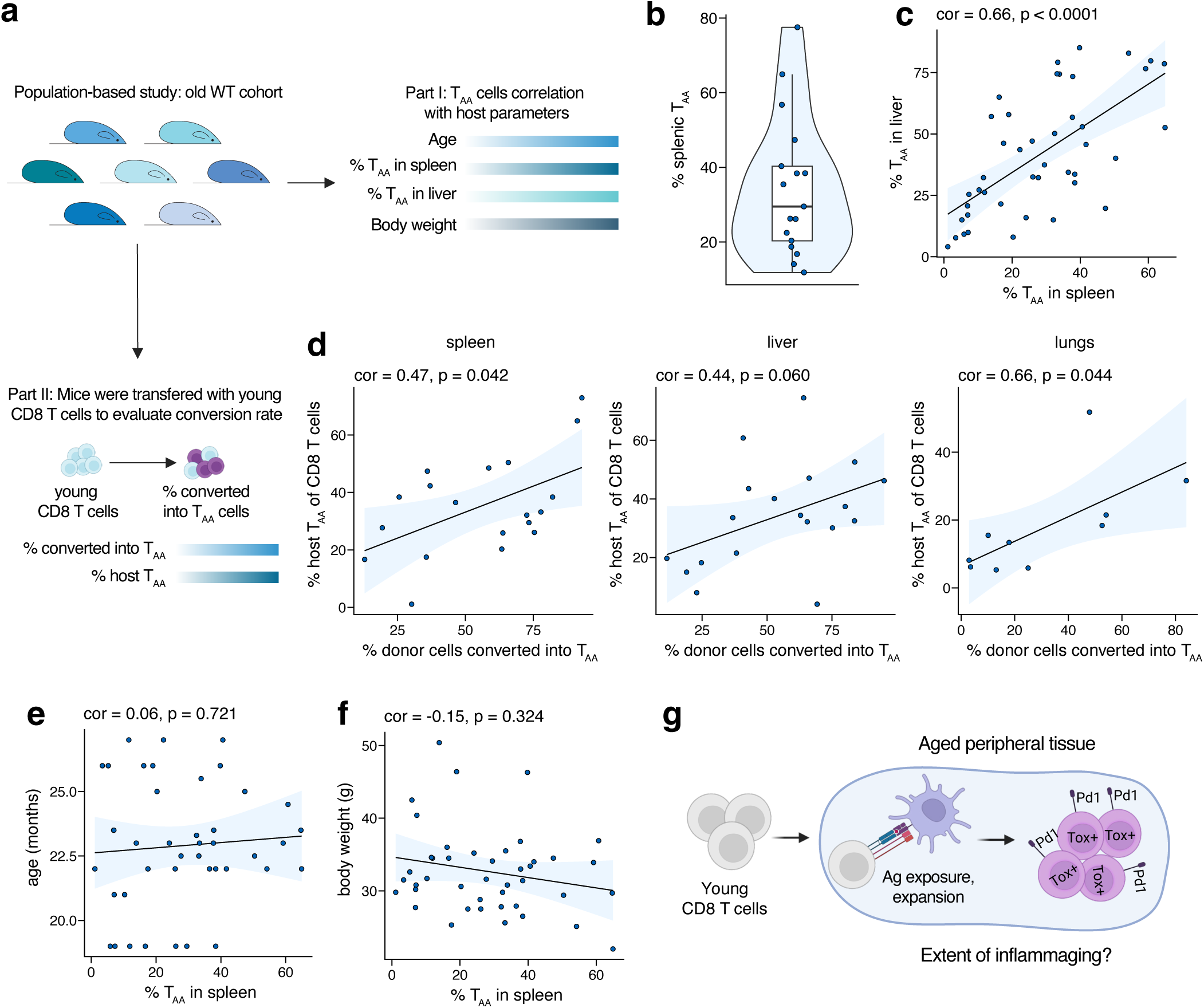
T_AA_ cell development and maintenance are regulated by environmental factors. **a.** Experiment design. **b.** Violin plot showing the variability of T_AA_ load in the spleens of old mice. **c.** Correlation between the frequency of T_AA_ cells in spleen and liver. **d.** Correlation between the frequency of host T_AA_ cells and the frequency of young donor CD8 T cells that de novo converted into T_AA_ cells in various tissues after transfer. **e-f**. Correlation between T_AA_ cell frequency and age (**e**) or body weight (**f**). **g.** Summary schematic. (**c-e**) Spearman’s rank correlation coeffecient. Exact group sizes are available in methods.

We next considered whether this regulation of T_AA_ cell abundance reflects static maintenance of existing cells, or if *de novo* generated cells exhibit a similar pattern. To explore this, we performed adoptive transfer experiments outlined in Fig. 1a using a larger cohort. Strikingly, the percentage of young donor CD8⁺ T cells that acquired the T_AA_ phenotype after transfer strongly correlated with the abundance of endogenous T_AA_ cells in the aged host (Fig. 2d), suggesting that the rate of early T_AA_ cell phenotype acquisition is also driven by environmental cues. Notably, among aged mice (>18 months old), T_AA_ cell abundance did not correlate with chronological age (Fig. 2e), arguing against a simple progressive accumulation over time. Correlation with body weight was modest and negative (Fig. 2f). Together, these findings indicate that the aged environment governs both the initiation and the extent of the T_AA_ program, shaping its magnitude and persistence across multiple tissues (Fig. 2g).

### Systemic chronic low-grade inflammation accelerates CD8^+^ T cell aging

In mice, inflammaging is a well-characterized hallmark of aging, defined by chronically elevated levels of pro-inflammatory cytokines such as IL-1β, IL-27, IL-6, and TNFα^5,21^. Given our findings implicating the aged environment in promoting T_AA_ cell development, we hypothesized that chronic low-grade inflammation may be sufficient to drive this process. Notably, previous models of accelerated immune aging have primarily focused on cell-intrinsic defects within T cells, such as impaired mitochondrial function^12^, dysregulated apoptosis^13^, or defective DNA repair^11^. To complement these findings, we next aimed to evaluate how T cell-extrinsic systemic low-grade inflammation shapes a CD8^+^ T cell compartment.

To test this, we used CRISPR-Cas9 gene editing to delete the AU-rich response element in the 3’ region of TNF mRNA to generate the TNF^Δ69AU/+^ mouse model (Fig. 3a, Supplementary Fig. 2a). This mutation would be expected to increase TNF mRNA stability, resulting in enhanced TNFα protein production from a single transcript^22^. However, as the manipulation did not further modify the TNF locus, a single copy of the mutated TNF locus did not lead to overt pathology like ileitis, contrasting with a previous model of TNF^ΔARE/+^ mice that had deleted the same AU-rich response element, while adding additional nucleotides associated with the targeting vector^22,23^. TNF^ΔARE/+^ mice had greatly elevated TNFα levels at the ileum, but TNF^Δ69AU/+^ mice did not (Supplementary Fig. 2b). TNF^ΔARE/+^ mice failed to gain weight by 8 weeks of age relative to littermate WT controls, but TNF^Δ69AU/+^ mice did not (Supplementary Fig. 2c), as the ileum of TNF^Δ69AU/+^ mice maintained a histological appearance similar to WT littermates (Supplementary Fig. 2d). Accordingly, fecal lipocalin levels in TNF^Δ69AU/+^ mice were not significantly elevated, whereas they were strongly increased in TNF^ΔARE/+^ mice (Supplementary Fig. 2e). Circulating TNFα levels were moderately elevated in TNF^Δ69AU/+^ mice (Fig. 3b). Canonical inflammaging-associated cytokines – IL-1β, IL-6, CCL2, and CXCL10^21^ – showed a trend toward increased levels, though these did not reach statistical significance (Fig. 3c).

**Fig. 3:**
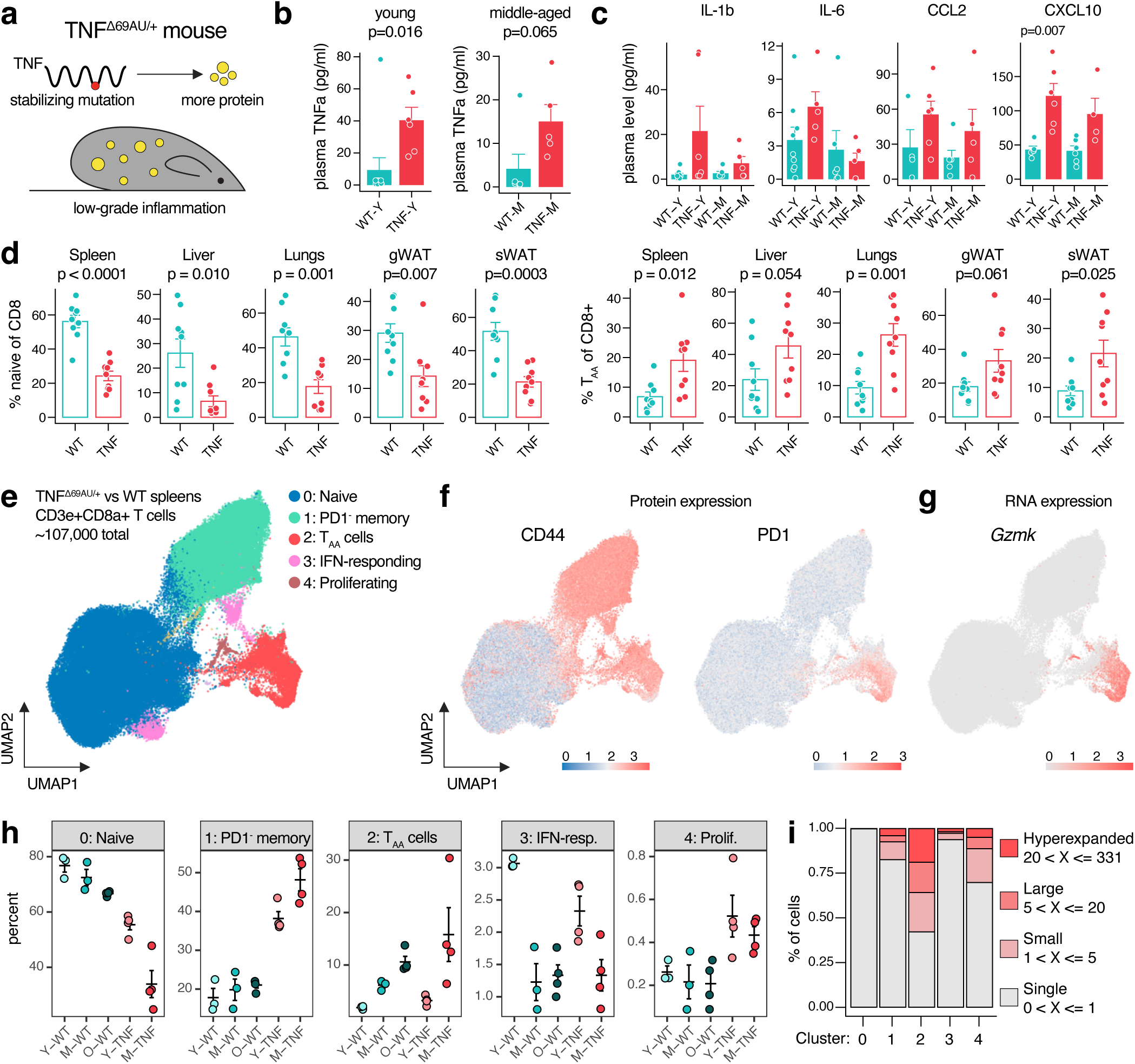
T_AA_ cells arise in response to low-grade chronic inflammation. **a**. Schematic of the TNF^Δ69AU/+^ mouse model. **b-c**. Circulating levels of TNFa (**b**) and selected cytokines (**c**) in young (5-8 weeks old) and middle-aged (13-16 months old) WT and TNF^Δ69AU/+^ mice. **d**. Frequency of naive and T_AA_ cells in various tissues of middle-aged WT and TNF^Δ69AU/+^ mice. **e**. UMAP plot of CD8 T cells isolated from the spleens of WT and TNF^Δ69AU/+^ mice. **f**. UMAP plot showing CD44 and PD1 protein expression. **g**. UMAP plot showing *Gzmk* transcript expression. **h**. Frequncy of corresponding clusters relative to the total number of sequenced CD8 T cells. Each dot represents an independent biological replicate. **i**. Relative clonality of major CD8 T cell clusters. Unpaired two-sample Student’s t-test **(b-d)**. Data are mean ± s.e.m. for (**b-d, h**). Exact group sizes are provided in methods.

Strikingly, TNF^Δ69AU/+^ mice displayed a pronounced phenotype of accelerated CD8^+^ T cell aging. In middle-aged animals (12-13 months old), we observed a marked reduction in naïve CD8^+^ T cells compared to age-matched WT controls (Fig. 3d). This was accompanied by an accumulation of PD1^−^ memory and PD1^+^TOX^+^ CD8^+^ T cells exhibiting the age-associated T_AA_ phenotype across multiple tissues (Fig. 3d, Supplementary Fig. 2c). Notably, some of these alterations were already detectable in young mice (Supplementary Fig. 2d-f). However, their overall profile still largely resembled WT controls, suggesting that the observed differences result from accelerated aging rather than a developmental defect.

To confirm that CD8^+^ T cells from TNF^Δ69AU/+^ mice reflect an aged phenotype and did not acquire an alternative cellular state, we performed single-cell transcriptomic analysis of splenic CD8^+^ T cells from TNF^Δ69AU/+^ and age-matched WT littermates (Fig. 3e, Supplementary Fig. 3a). Transcriptionally defined populations aligned well with surface marker expression of PD1 and CD44 (Fig. 3e-g). Cells from TNF^Δ69AU/+^ and WT mice clustered together and showed concordance with flow cytometric quantification (Fig. 3h, Supplementary Fig. 3b-d). Transcriptomic data corroborated the age-related trend: naïve cells were significantly depleted in young TNF^Δ69AU/+^ mice, while PD1^−^ memory and T_AA_ cells accumulated more rapidly than in WT controls (Fig. 3h). Intriguingly, TNF^Δ69AU/+^ mice also harbored a higher proportion of proliferating cells (Fig. 3h). As previously observed, *Gzmk*-expressing cells were restricted to the PD1^+^CD44^+^ T_AA_ cell cluster, which was also the most clonally expanded (Fig. 3g, i). Collectively, these data demonstrate that CD8^+^ T cells from TNF^Δ69AU/+^ mice closely resemble those from aged WT animals, indicating that chronic low-grade inflammation is sufficient to accelerate CD8^+^ T cell aging.

### T_AA_ cells are heterogeneous and include a progenitor population

To gain deeper insight into T_AA_ cell heterogeneity, we performed a focused analysis of the splenic PD1^+^CD44^+^ cluster (Fig. 4a, Supplementary Fig. 4a-b). It revealed multiple distinct subsets, including minor populations previously undetected due to low cell numbers in single-cell experiments. For example, we identified *Foxp3*-expressing CD8^+^ regulatory cells that were increased in aged mice (Fig. 4b). We also observed a variable but overall increasing trend in the frequency of *Klrg1*⁺ effector cells and *P2rx7*⁺ CD8⁺ T cells with age (Fig. 4b).

**Fig. 4:**
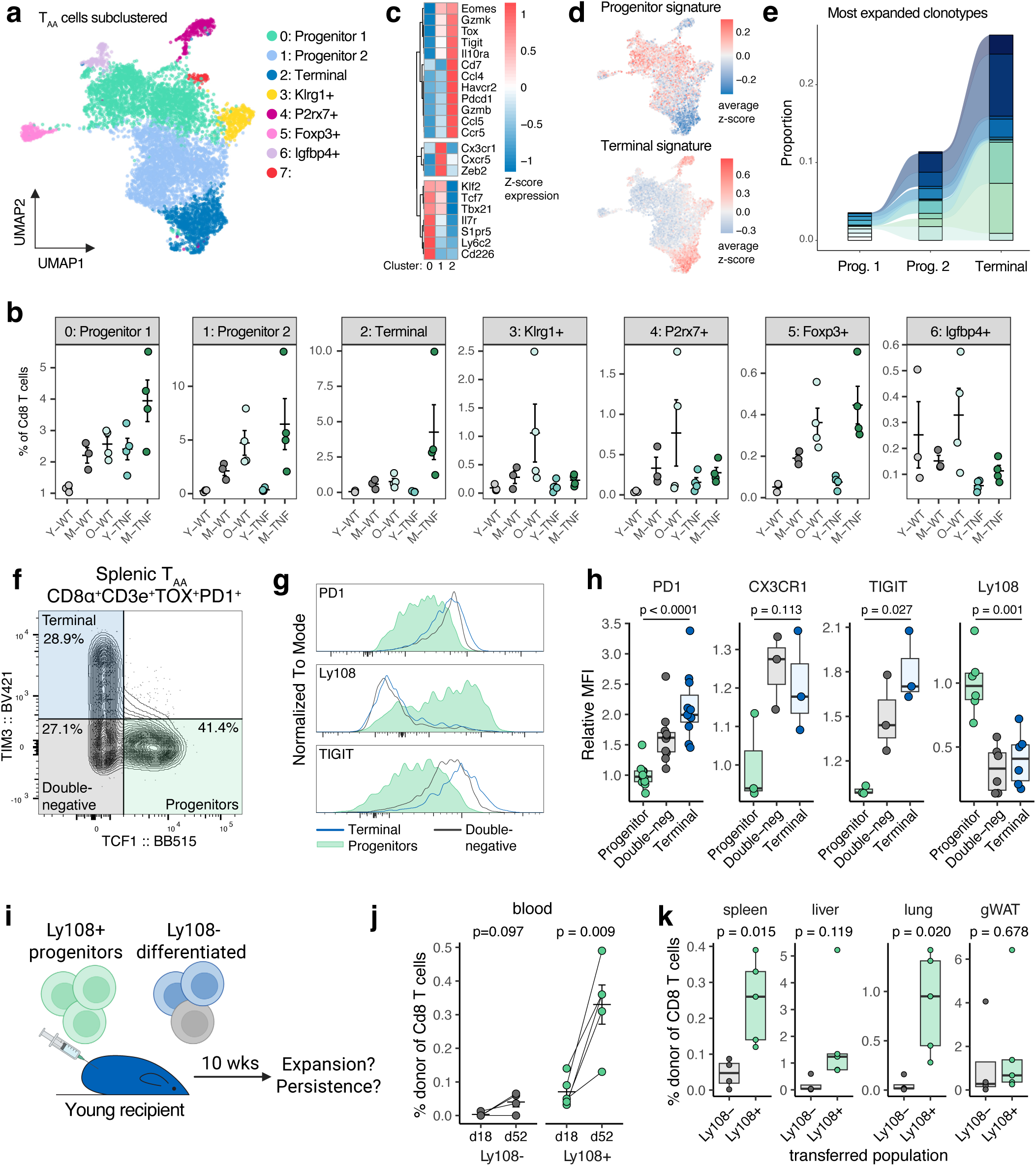
T_AA_ cells contain a functional progenitor subpopulation. **a**. UMAP plot of CD44+PD1+ CD8 T cells (cluster 2 from Fig. 3e). **b**. Heatmap of selected gene expression in clusters 0, 1, and 2. **c**. UMAP plot showing average z-score of progenitor and terminal exhausted CD8 T cell signatures derived from GSE149876. **d**. Alluvial plot tracking the size of non-shared and shared TCR clones across clusters 0, 1, and 2. **e**. Frequency of corresponding clusters relative to the total number of sequenced CD8 T cells. Each dot represents an independent biological replicate. **f**. Definition of T_AA_ subpopulations based on TIM3 and TCF1 protein expression. **g-h**. Protein expression of selected markers in splenic T_AA_cell subpopulations. **i**. Transfer experiment design. **j**. Frequency of donor cells in the recipient’s circulation at indicated time points post-transfer. **k**. Frequency of donor cells in recipient tissues ∼2 months post-transfer. Unpaired two-sample Student’s t-test (**h, k**). Paired two-sample Student’s t-test (**j**). Data are mean ± s.e.m. for (**b, j**). Exact group sizes are provided in methods.

The majority of PD1⁺CD44⁺ CD8⁺ T cells were contained within three major clusters. Clusters 0 and 1 expressed classical progenitor exhausted cell markers such as *Tcf7*^24^ and *Klf2*^25^, while Cluster 2 represented a terminally exhausted population marked by high expression of inhibitory receptors^26^ (*Havcr2, Tigit, Pdcd1*) and effector molecules^26^ (*Gzmk, Gzmb, Ccl5*) (Fig. 4c, Supplementary Fig. 4a-b). Furthermore, we found that the gene expression profiles of the clusters aligned closely with previously described progenitor and terminal exhaustion states from chronic infection models^27^ (Fig. 4d). TCR analysis revealed that terminal cells were the most clonally expanded of the three subsets, and the substantial TCR overlap across clusters indicates that these are likely the part of a common developmental trajectory (Fig. 4e). Relative to the total CD8⁺ T cell population, both progenitor clusters increased with age, while terminally differentiated cells showed the most pronounced expansion in middle-aged TNF^Δ69AU/+^ mice (Fig. 4b).

CD8^+^ T cell exhaustion is actively studied in the context of chronic viral infections and cancer, leading to the identification of reliable markers of progenitor and terminally differentiated cells^27^. At the protein level, progenitors are commonly marked by TCF-1 expression^24^, while TIM3 marks terminally exhausted cells^28,29^, consistent with our transcriptomic findings (Fig. 4b, Supplementary Fig. 4a-b). A double-negative population likely represents intermediate differentiation states. Using this established panel, we profiled T_AA_ cells in the spleen and identified three distinct subpopulations by flow cytometry, recapitulating standard gating strategies (Fig. 4f). As expected, progenitor cells also expressed high levels of Ly108 – a surface marker frequently used to sort this subset^27^ – and exhibited significantly lower levels of inhibitory receptors such as PD1 and TIGIT (Fig. 4g-h).

Having identified a progenitor-like population based on transcriptomic and protein expression profiles, we next asked whether this subset could sustain and repopulate the T_AA_ compartment. To test this, we conducted a transfer experiment (Fig. 4i). Our prior work showed that T_AA_ cells transferred into young mice persist and maintain their aged phenotype, providing a model to study their maintenance^5^. We sorted Ly108^+^ cells (enriched for progenitors, Supplementary Fig. 4c) and Ly108^−^ cells (mostly differentiated T_AA_ cells Supplementary Fig. 4c), transferred them into young recipients, and tracked donor-derived cells in the circulation. Approximately two months post-transfer, we observed robust expansion of the T_AA_ population only in mice that received Ly108^+^ progenitors (Fig. 4j), which also exhibited higher abundance in tissues (Fig. 4k). These findings suggest that T_AA_ cells harbor a functional progenitor subset capable of expansion and tissue repopulation, whereas more differentiated T_AA_ cells lack this capacity.

### Age-associated *Gzmk*^+^ CD8^+^ T_AA_ cells are characterized by a CD226^−^PD1^+^CD44^+^ phenotype

While all three major subsets of T_AA_ cells were accumulating with age (Fig. 4e), we observed a clear age-associated shift in the composition of the progenitor compartment (Fig. 5a-c, Supplementary Fig. 5a-b). Although rare in absolute numbers, Progenitor 1 cells were already present in young mice and comprised the majority of T_AA_ cells in the young spleen (Fig. 5a-c). With age, these cells became scarce, and Progenitor 2 cells emerged as the dominant subset in the spleen (Fig. 5a-c). Terminally differentiated cells also increased in middle-aged and old mice, with the most pronounced expansion in middle-aged TNF^Δ69AU/+^ mice (Fig. 5a-c).

**Fig. 5:**
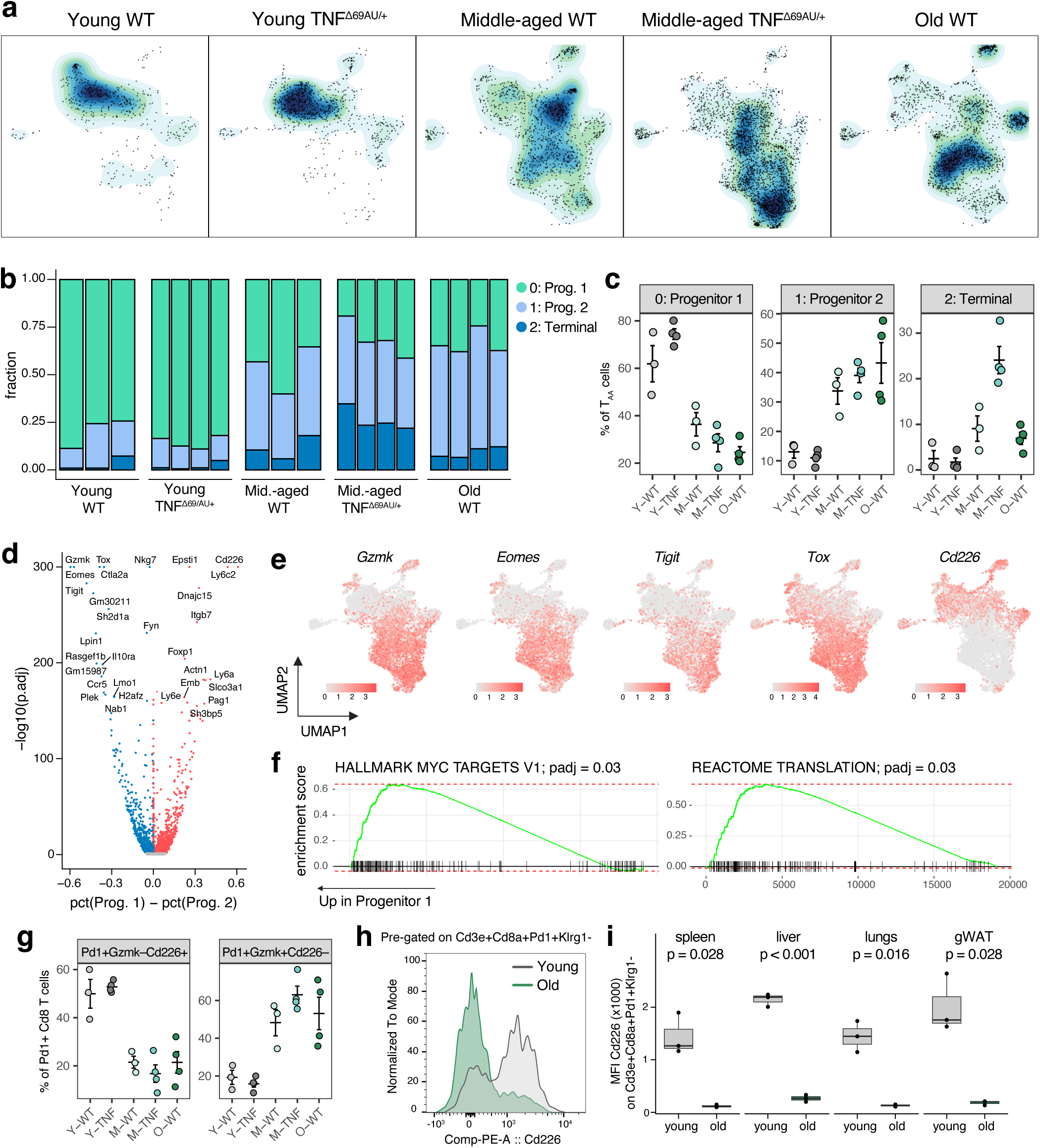
CD226^−^GZMK^+^ is an age-dependent subpopulation of PD1^+^ CD8^+^ T cells. **a**. UMAP density plots showing the distribution of T_AA_ cells across ages and genotypes. **b**. Bar plot illustrating the composition of major T_AA_ subset. Each bar represents an individual mouse. **c**. Frequency of corresponding clusters relative to the total number of T_AA_ cells. Each dot represents an independent biological replicate. **d**. Volcano plot summarizing differential expression analysis between Progenitor 1 and Progenitor 2 clusters. **e**. UMAP plots showing the expression of selected transcripts. **f.** Gene set enrichment curves for pathways enriched in Progenitor 1 population. **g**. Frequency of corresponding T_AA_ cell subsets relative to the total number of T_AA_ cells. Each dot represents an independent biological replicate. Cells were defined as positive for Gzmk and Cd226 if at least one transcript count was detected. **h.** Protein expression of CD226 in splenic T_AA_cell subpopulations. **i.** Quantified CD226 protein expression on T_AA_ cells from different tissues of old and young mice. Unpaired two-sample Student’s t-test (**i**). Data are mean ± s.e.m. for (**c, g**). Exact group sizes are provided in methods.

Strikingly, while investigating molecular differences between the progenitor subsets, we observed that Progenitor 1 largely lacked expression of *Gzmk* – the major transcriptional marker of T_AA_ cells^5^ (Fig. 5d-e, Supplementary Fig. 5c). Other markers such as *Eomes* and *Tox* were also significantly down-regulated in Progenitor 1 cells (Fig. 5d-e). In contrast, Progenitor 1 cells were enriched in MYC targets and translation-related genes, consistent with MYC’s role as an early activation marker and suggesting these are the least differentiated cells^30^ (Fig. 5f). Among other top differentially expressed genes was *Cd226*, encoding the co-stimulatory receptor DNAM-1 (Fig. 5d-e). In young mice (both WT and TNF^Δ69AU/+^), most T_AA_ cells expressed *Cd226* but lacked *Gzmk* (Fig. 5g, Supplementary Fig. 5d). This pattern reversed with age, as older mice showed increased *Gzmk* and reduced *Cd226* expression (Fig. 5g, Supplementary Fig. 5d). We validated these findings at the protein level using flow cytometry, demonstrating a similar age-related loss of CD226 on PD1^+^ CD8^+^ T cells in the spleen and other tissues (Fig. 5h-i). Our results demonstrate that CD226⁻PD1⁺CD44⁺ CD8^+^ T cells represent an age-dependent *Gzmk*⁺ CD8^+^ T_AA_ cell population that is transcriptionally distinct from CD226^+^PD1⁺CD44⁺ CD8^+^ T cells observed in some young tissues. These findings significantly refine our understanding of CD8^+^ T cell aging and T_AA_ cells in particular.

### Adipose tissue harbors T_AA_ cell progenitors

Given the presence of distinct progenitor and terminal subpopulations of T_AA_ cells in the spleen, we next investigated how this heterogeneity is represented across different tissues. We discovered a markedly tissue-specific distribution of progenitor and terminal T_AA_ cells among organs. While the spleen exhibited a mixture of all three subpopulations (Fig. 4e), the liver and lungs were predominantly populated by TCF-1-negative, more differentiated terminal T_AA_ cells, with progenitor cells nearly absent (Fig. 6a-b). In contrast, progenitor T_AA_ cells were highly enriched in gonadal adipose tissue (Fig. 6a-b). Together with our previous data demonstrating the importance of peripheral tissues and progenitor T_AA_ cells in T_AA_ cell development and maintenance, these findings strongly suggest that adipose tissue might serve as a critical niche.

**Fig. 6:**
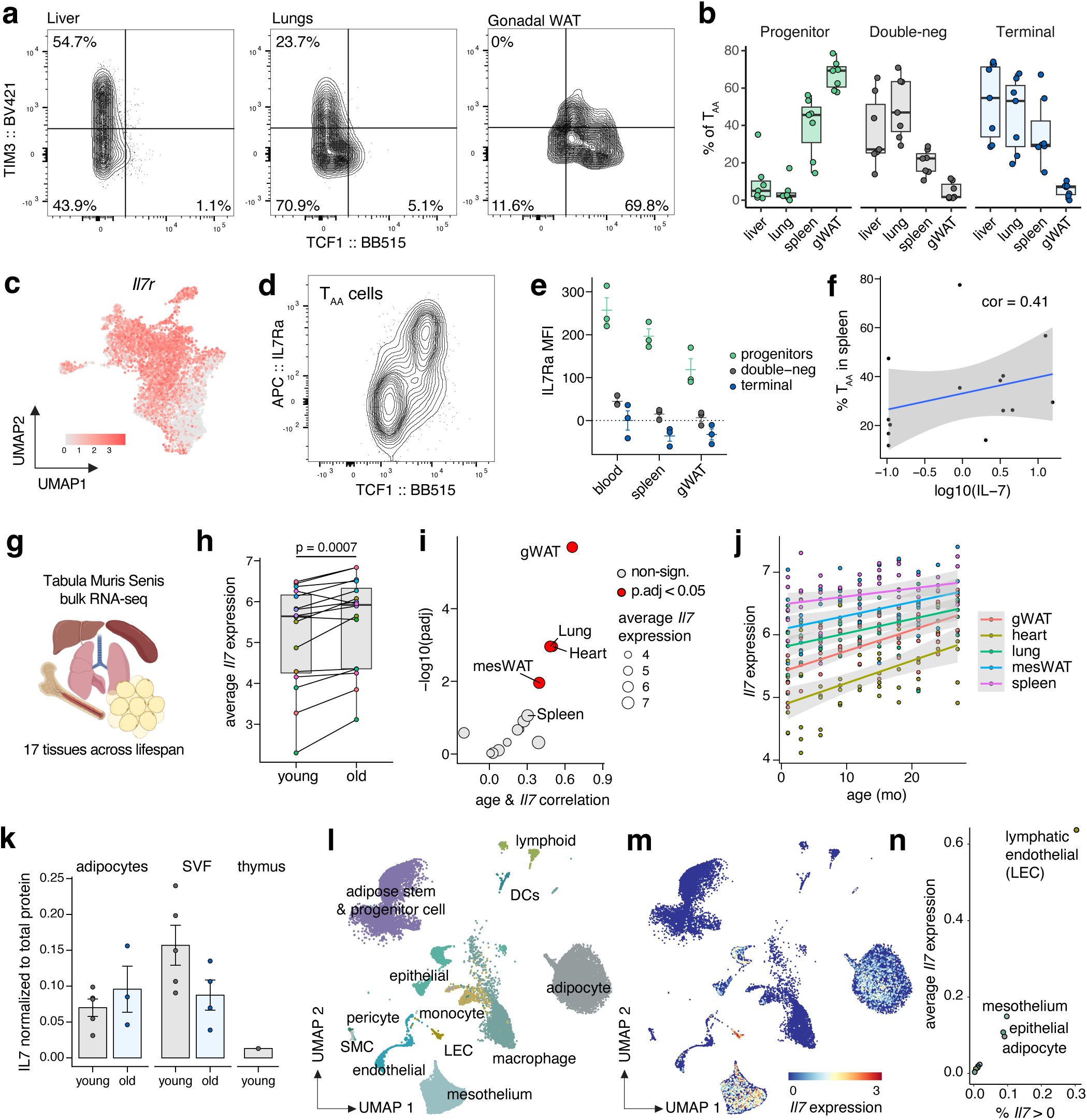
T_AA_ progenitors are enriched in adipose tissue. **a.** TIM3 and TCF1 expression on T_AA_ cells from different tissues. **b**. Frequency of T_AA_ subpopulations in various tissues. **c**. UMAP plot showing *Il7r* transcript expression in T_AA_ cells. **d**. Protein expression of IL7Rα on splenic T_AA_ cells. **e**. Quantification of IL7Rα expression on T_AA_ cells from different tissues. **f**. Correlation between plasma IL7 levels and frequency of T_AA_ cells among total CD8 T cells in spleen. **g**. Tabula Muris Senis data summary schematic. **h**. Average *Il7* expression in different tissues from old and young mice. **i**. Correlation between *Il7* expression and mouse age versus corresponding adjusted p-value. Each dot represents one tissue. **j**. Correlation between *Il7* expression and mouse age for selected tissues. **k**. IL7 protein levels normalized to total protein across different tissues. **l**. UMAP plot of snRNA-seq data from gonadal adipose tissue of chow-fed young mice (from Emont et al.) **m**. UMAP plot showing *Il7* expression. **n**. Summary of *Il7* expression by cell type. Spearman’s rank correlation coeffecient (**f**). Paired two-sample Student’s t-test (**h**) and Pearson’s correlation with Benjamini-Hochberg correction (**j**) on average normalized data (see Methods). Data are mean ± s.e.m. for (**e, k**). Exact group sizes are available in methods.

Previous studies have reported that the scavenger receptor CD36 is upregulated on T cells in adipose tissue^31^ and can promote an exhausted phenotype^32^. However, we did not observe any difference in T_AA_ cell abundance between aged CD36 knockout and wild-type mice (Supplementary Fig. 6a), suggesting that CD36 is not required for T_AA_ cell development. To explore other potential regulators of T_AA_ progenitors, we analyzed differentially expressed genes between progenitor and terminal T_AA_ subsets (Supplementary Fig. 6b). Among the most upregulated genes in progenitors was *Il7r*, which encodes the IL-7 receptor (Fig. 6c, Supplementary Fig. 6b-c). We also confirmed that it was highly co-expressed with TCF-1 at the protein level (Fig. 6d-e). Furthermore, we observed a positive correlation between the abundance of T_AA_ cells in individual mice and their circulating IL-7 levels (Fig. 6f).

To identify the source of IL-7, we turned to the Tabula Muris Senis (TMS) dataset^33^, which includes bulk RNA-seq data from multiple tissues across the mouse lifespan (Fig. 6g). We first asked whether IL-7 expression increases with age. Comparing average *Il7* expression in various tissues between young and old mice revealed a consistent, global upregulation in aged animals (Fig. 6h). In nearly all tissues examined, *Il7* expression positively correlated with age, with statistically significant increases in mesenteric white adipose tissue, lung, heart, and gonadal white adipose tissue (Fig. 6i-j). Given that transcriptional changes do not always reflect protein abundance, we next measured IL-7 protein in adipose tissue using ELISA on tissue lysates (Fig. 6k). When normalized to total protein, IL-7 was more abundant in aged adipose tissue than in young thymus, where it is known to be essential for T cell development^34^ (Fig. 6k). IL-7 protein was detectable in both adipocyte and stromal vascular fractions (SVF) (Fig. 6k). To identify specific IL-7-producing cells, we analyzed publicly available single-nucleus RNA-seq data from adipose tissue of young lean mice^35^ (Fig. 6l). Consistent with our ELISA results, *Il7*-expressing cells were found among adipocytes (Fig. 6m). Within the SVF, lymphatic endothelial cells emerged as the primary source of *Il7* expression (Fig. 6m-n).

Together with recent findings highlighting the role of adipose tissue as a reservoir for memory CD8^+^ T cells^36^, our data suggest that this tissue plays a pivotal role in the development of T_AA_ cells in aged mice. Thus, we next tested whether this tissue alone is sufficient to drive the generation of T_AA_ cells.

### Old adipose tissue is sufficient for T_AA_ cell development

To test if old adipose tissue alone is sufficient for the development of T_AA_ cells, we performed adipose tissue transplants (Fig. 7a). We took equally sized pieces of gonadal fat pads from young and old CD45.2^+^ mice and subcutaneously implanted them on the back scruff of young CD45.1^+^ recipients^37,38^. A month later, CD8^+^ T cells in the recipient tissues were analyzed.

**Fig. 7:**
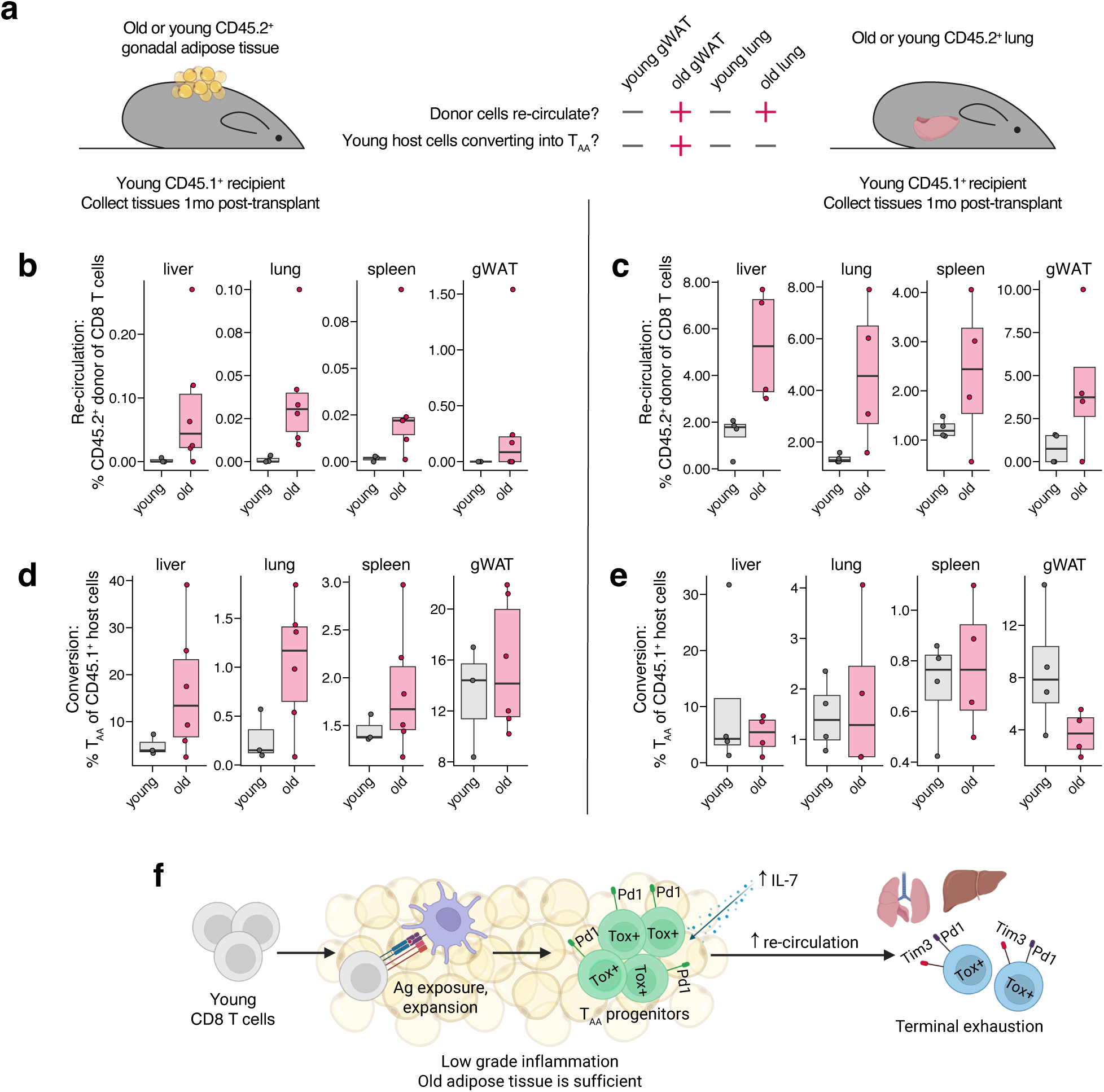
Old adipose tissue is sufficient for the development of T_AA_ cells. **a.** Experimental design of the transplant experiments. **b-c.** Frequency of donor CD8 T cells detected in host tissues, representing CD8⁺ T cells that exited the adipose (**b**) or lung (**c**) transplant and seeded recipient tissues. **d-e.** Frequency of PD1^+^TOX^+^ recipient CD8^+^ T cells, indicating the conversion of young host cells into T_AA_ cells following adipose (**d**) or lung (**e**) transplant. **f.** Summary schematic.

Our model of T_AA_ cell generation assumed that T_AA_ cells originating from progenitor populations exit the fat and populate peripheral tissues. However, in young mice, CD8^+^ T cells in adipose tissue are thought to be highly resident with no evidence of recirculation, as previously shown by parabiosis experiments^36^. Therefore, our first question was whether CD8^+^ T cells from old adipose tissue could exit the transplant. Encouragingly, we found CD45.2^+^ donor CD8^+^ T cells originating from the adipose transplant in all profiled tissues of the CD45.1^+^ recipients who received old transplants (Fig. 7b). In contrast, no cells exited from young fat transplants, consistent with previous reports. These results indicate that old adipose tissue becomes more circulatory, contributing to increased systemic abundance of T_AA_ cells across old tissues. To isolate the contribution of the donor adipose tissue, we repeated the experiment using AdiponectinCre/+ RosaDTAfl/+ fatless young recipients that lack mature adipocytes and observed a similar pattern (Supplementary Fig. 6d). As a control, we also performed lung transplants following the same study design (Fig. 7a). Old lungs primarily contain terminally differentiated T_AA_ cells, so we expected they would not drive T_AA_ cell re-population as effectively as adipose tissue (Fig. 4i-k). However, we observed a similar increase in the overall trafficking of CD8^+^ T cells from the transplanted old lungs compared to young lungs (Fig. 7c).

We next examined host CD8^+^ T cells to assess whether the transplanted tissues influenced their phenotype. All recipient mice were young, so almost no native CD45.1^+^ T_AA_ cells were expected in their tissues. However, a month after receiving old adipose tissue transplants, host CD8^+^ T cells began acquiring a T_AA_ cell phenotype (Fig. 7d). While the increase was mild, this observation suggests that transplanting old fat into young mice can influence the CD8^+^ T cell landscape in a systemic manner. Importantly, neither young fat transplants nor old and young lung transplants promoted this shift, suggesting that this feature is specific to aged adipose tissues (Fig. 7d, e).

Overall, our data lead us to propose the following T_AA_ cell development and maintenance model (Fig. 7f). Following antigen exposure in old tissues outside the spleen and lymph nodes, a progenitor T_AA_ cell population is formed in the old gonadal fat. These cells respond to IL-7, which increases systemically with age and is abundant in the adipose tissue. Furthermore, our results demonstrate that old adipose tissue allows for the recirculation of CD8^+^ T cells, leading to the systemic distribution of T_AA_ cells in old mice. The progenitor population can thus expand and populate peripheral tissues, where T_AA_ cells are predominantly terminally differentiated, which suggests this is the endpoint of their lifespan.

## DISCUSSION

In this study, we demonstrate that a mildly pro-inflammatory aged peripheral environment can systemically reshape the CD8⁺ T cell compartment and drive the development of age-associated CD8⁺ T cells (T_AA_ cells). T_AA_ cell development occurs in an antigen-dependent manner within non-lymphoid peripheral tissues and is driven by local environmental cues. Within the T_AA_ population, we identified a functional progenitor subset capable of seeding multiple tissues throughout the body. These progenitor cells are particularly enriched in the white adipose tissue of aged mice, implicating this tissue as a key niche for T_AA_ cell maintenance. Supporting this, we show using a transplantation model that aged – but not young – adipose tissue is sufficient to induce T_AA_ cell development.

These findings contribute to a growing body of evidence linking aging and immune system dysfunction. While multiple studies have shown that an abnormal immune system can accelerate the onset of age-associated phenotypes^9,10^ – including the establishment of chronic low-grade inflammation, or “inflammaging” – our results demonstrate that the reverse is also true: the aged, inflamed environment itself can reprogram immune cells. CD8⁺ T cells, in particular, are highly responsive to environmental signals. When exposed to persistent, low-grade inflammation – whether due to natural aging or genetic perturbation – a substantial fraction of CD8⁺ T cells acquires an exhausted, age-associated phenotype. This transition is antigen-driven and occurs in peripheral tissues, enabled in part by altered trafficking patterns that emerge with age. For example, aged adipose tissue becomes more permissive to T cell infiltration and egress, potentially facilitating T cell reprogramming through sustained exposure to local antigens and inflammatory cues. Further investigation is needed to understand how age-related changes in immune cell trafficking affect immune function and to identify the specific molecular drivers of these alterations. In the context of T_AA_ cell development, we ruled out previously implicated in abnormal CD8⁺ T cell migration chemokine CXCL13^19^ and T_AA_ cell marker CD49d^5^.

T_AA_ cells were originally described as a presumably homogeneous population characterized by *Gzmk* transcript expression and co-expression of PD1 and TOX at the protein level^5^. However, in this study, we reveal greater heterogeneity within these cells. Specifically, we identify progenitor subsets of T_AA_ cells marked by expression of TCF-1, representing a less differentiated T_AA_ cell state capable of populating various tissues. Within these progenitors, we further distinguish two major subsets. Progenitor 1 cells are the least differentiated PD1⁺ cells that are already present in young mice. Notably, these cells do not express *Gzmk*, a gene that becomes upregulated only at the Progenitor 2 stage, which expands dramatically with age. These two subsets can be distinguished by surface expression of CD226, which marks Progenitor 1 cells and is largely absent in aged mice. The identification of CD226 as a marker now enables the separation of *Gzmk*⁺ and *Gzmk*⁻ subpopulations within the broader PD1⁺ T_AA_ cell pool. Overall, our findings provide a significantly more nuanced understanding of the population structure and age-related dynamics of CD8⁺ T cells.

The identification of a progenitor subset enabled us to pinpoint peripheral tissues that influence T_AA_ cell development. We found that T_AA_ cell progenitors are enriched in white adipose tissue, while in tissues such as the liver and lungs, T_AA_ cells are predominantly terminally differentiated. This suggests a model in which aged adipose tissue serves as a key niche for T_AA_ cell development. Within this niche, progenitors may either arise through local conversion or be maintained in an undifferentiated state through the provision of survival and maintenance signals, such as IL-7. These progenitor cells can then recirculate and populate other peripheral tissues, where they undergo terminal differentiation. This model highlights an underappreciated role for adipose tissue in regulating CD8⁺ T cell homeostasis. Adipose tissue has already been identified as an important reservoir for memory CD8⁺ T cells^36^, but its influence on T cell aging and differentiation remains poorly understood. Thus, perturbations in adipose tissue, such as those induced by obesity, may have profound immunological consequences. Notably, *Gzmk*⁺ CD8⁺ T cells have been shown to expand in high-fat diet-induced obesity models and persist even after weight loss^39^. These findings underscore the critical importance of the tissue environment – particularly its metabolic and inflammatory status – in shaping CD8⁺ T cell fate and function.

Together, these findings have important implications for the design of immune-based therapies. Strategies that rely on the introduction of functional CD8⁺ T cells into aged or inflamed hosts – such as adoptive cell therapies for cancer or thymic rejuvenation approaches – may be compromised if the host environment rapidly reprograms the infused cells into dysfunctional or exhausted states. A deeper understanding of the mechanisms driving T_AA_ cell development, including the role of tissue-specific antigens, altered trafficking, and local antigen presentation, will be critical for improving the durability and efficacy of such interventions.

## Authors contributions

I.S. and M.N.A. conceived and designed the project and wrote the original draft. I.S. and C.J.R.H. performed the experiments and data analysis. H.S.R. and G.J.R. designed, developed, and characterized the TNF^Δ69AU/+^ mouse. M.K. assisted with computational analysis. R.L.M., S.C.G., K.M., D.W.P. and D.K. assisted with transplantation experiments. S.H. and T.E. helped with LCMV infections. V.V., J.K., D.A.M., C.G.H., B.E. assisted with experiments and mouse handling. I.S. prepared figures. M.N.A. and G.J.R. acquired the funding. All authors reviewed and edited the manuscript.

## Acknowledgments

The study was supported by funding from the Aging Biology Foundation to MNA., NIH grants R01AI168044, R37AI049653, and U01AI163064 to GJR and F30CA281124 to RLM. HSR is funded by NIH grant T32DK007130.

We thank Dorjan Brinja, Dr. Erica Lantelme, Pascaline Akitani and Flow Cytometry & Fluorescence Activated Cell Sorting Core at Washington University for their help with cell sorting and flow cytometry. We thank Molly Keppel and Siteman Flow Cytometry Core at Washington University for assistance with cell sorting. We thank Dr. Sheila Stewart for providing aged mice.

Additional support was provided by the Digestive Diseases Research Core Center (P30DK052574) and the Bursky Center for Human Immunology and Immunotherapy at Washington University. The TNF^ΔARE/+^ mouse line was provided by Fabio Cominelli through the Cleveland Digestive Disease Research Core Center (P30DK097948) to GJR. We thank the Genome Engineering & Stem Cell Center (GESC@MGI) at the Washington University for CRISPR reagents design and validation services in the generation of a new mouse strain. We are grateful for the assistance of Mike White, Mary Wohltmann, and Galina Fedotova for their assistance in the generation and genotyping of TNF^Δ69AU/+^ mice, and for their care and maintenance. We extend thanks to Emma Erlich and Michael Strickland for helpful suggestions.

We thank the Genome Technology Access Center at the McDonnell Genome Institute at Washington University School of Medicine for help with library generation, sequencing, and analysis. The Center is partially supported by NCI Cancer Center Support Grant #P30 CA91842 to the Siteman Cancer Center from the National Center for Research Resources (NCRR), a component of the National Institutes of Health (NIH), and NIH Roadmap for Medical Research. This publication is solely the responsibility of the authors and does not necessarily represent the official view of NCRR or NIH.

## METHODS

### Mice

Wild-type (WT) C57BL/6J mice were obtained from the Jackson Laboratory (strain #000664) and bred and aged at Washington University in St. Louis. Some aged mice were acquired from the National Institute on Aging (USA). B6.SJL-PtprcaPepcb/BoyJ (CD45.1, strain #002014) and B6.129S1-Cd36tm1Mfe/J (CD36KO, strain #019006) were purchased from the Jackson Laboratory. OT-1/RAG2 KO mice were purchased from Taconic (strain #2334) and bred to CD45.1 background. Adiponectin^Cre^ mice (The Jackson Laboratory, Bar Harbor, ME, USA, 028020) were bred with lox-stop-lox-Rosa26 diphtheria toxin (DTA) mice (the Jackson Laboratory, strain #010527, RRID: IMSR_JAX:028020) to generate AdiponectinCre/+ RosaDTAfl/+ fatless offspring that lack brown and white mature adipocytes. To compensate for the absence of adipose tissue and thermogenic capacity, all fatless mice and their littermate controls were housed at thermoneutrality (30°C) from birth until experimental procedures were initiated. The TNF^ΔARE/+^ mouse line was provided by Fabio Cominelli through the Cleveland Digestive Disease Research Core Center (P30DK097948) to GJR. All mice were maintained at Washington University in St. Louis in accordance with federal and university guidelines, following protocols approved by the Animal Studies Committee of Washington University in St. Louis. Age groups were categorized as old (greater than 18 months) and young (less than 6 months). Both male and female mice were included in the study.

### Generation of the TNF^Δ69AU/+^ mouse line

Targeting guides for CRISPR-Cas9-mediated excision of the AU-rich response element were designed at the Genome Engineering and Stem Cell Center of the McDonnell Genome Institute at Washington University. The targeted region and the chosen guides are shown in Supplementary Fig. 7a. Guides were selected based on fit to target, lack of presence of SNP in the guide, and minimal predicted off-target potential, using Doench 2016 scoring^40^ (Supplementary Fig. 7b). Guides complexed with Cas9 were electroporated into JM8.N4 mouse C57BL/6N embryonic stem cells. Then correctly targeted clones were injected into C57BL/6 blastocysts. F1 progenies were screened by PCR, next generation sequencing, and monitored for fecal lipocalin levels. Sequencing through the targeted locus in F1 or F2 heterozygous offspring revealed a precise removal of the targeted 69-bp region (Supplementary Fig. 7a), compared with a 70-bp excision in the previously generated TNF^ΔARE/+^ strain that targeted the same locus^22^ (Supplementary Fig. 7a), and no obvious off-target modifications. PCR across the targeted region, using the primers previously described^22,23^, accordingly led to a smaller band compared with WT, considering that the 69-bp was removed (Supplementary Fig. 7c). By contrast, whereas a 70-bp deletion was observed in the TNF^ΔARE/+^ mouse in the AU-rich response element, the targeting strategy used to generate the TNF^ΔARE/+^ mouse resulted in the addition of a 116-bp region in the 3’ of the TNF mRNA (Supplementary Fig. 7d), such that genotyping with the same primers leads to a larger, rather than smaller, PCR product (Supplementary Fig. 7c). To generate further progeny including WT littermate controls, WT C57BL/6 males were mated with female TNF^Δ69AU/+^ mice and the progeny used for the studied described. These studies were done in accordance with the Institutional Care and Use Committee approved protocol to GJR (22-0433).

### Adoptive CD8 T cell transfers

Bulk CD8^+^ T cells were enriched from the splenocytes using the EasySep Mouse CD8^+^ T Cell Isolation Kit (cat. #19853), following the manufacturer’s protocol. Two runs of the protocol were performed to ensure the purity. 0.5-2.5*10^6 donor CD8^+^ T cells in 100 µl PBS were injected intravenously via retroorbital injection into the isoflurane-anesthetized recipients.

To evaluate the proliferation rate post-transfer, donor cells were stained with CellTrace Far Red (Invitrogen, cat. #C34564) according to the manufacturer’s recommendations right before the injection. Briefly, isolated CD8^+^ T cells were resuspended in CellTrace Far Red diluted in PBS 1:1000 and incubated at 37°C for 20 minutes. Then, 5x amount of PBS containing 2% FBS and 1 mM EDTA was added, and cells were kept at 37°C for another 5 minutes and centrifuged at 500 g for 10 minutes.

In the FTY720 experiment (Fig. 1f), mice were injected i.p. daily with FTY720 (10 µg in 200 µl sterile PBS, Sigma-Ardrich cat. #SML0700) or sterile PBS. Anti-CXCL13 (500ug/mouse, clone 5378, kindly supplied by Vaccinex Inc.) and anti-CD49d (250ug/mouse, clone PS/2, BioXCell, cat. #BE0071) antibodies were injected every 3 days.

### Immune cell isolation

Spleens were collected, and splenocytes were isolated using mechanical dissociation. The cell suspension was filtered through the 70 µm strainer and pelleted by centrifugation at 500 g for 10 minutes at 4°C. Cells were resuspended in red blood cells (RBC) lysis buffer, incubated for 5 minutes at room temperature and washed once in FACS buffer (PBS containing 2% FBS and 1 mM EDTA).

Liver and lungs were minced into small pieces using scissors and incubated with shaking for 60 minutes at 37°C in a digestion solution containing collagenase D (Roche, cat. #11088882001) at a concentration of 0.1% by weight in RPMI 1640 (Gibco, cat. #21870-076) supplemented with 10% FBS and 1x PenStrep (Gibco, cat. #15140-122). After incubation, the samples were vortexed for 10 seconds and then passed through an 18-gauge needle to disrupt the tissue and facilitate cell release. The resulting cell suspension was filtered through a 70 µm strainer. Tissue debris was removed by centrifugation at 500 g for 30 minutes in a 30% Percoll (Cytiva, cat. #17544502) gradient at room temperature without using a brake. Cells were resuspended in RBC lysis buffer, incubated for 5 minutes at room temperature and washed once in FACS buffer.

Adipose tissue was minced into small pieces using scissors and incubated with shaking for 60 minutes at 37°C in a digestion solution containing collagenase D (Roche, cat. #11088882001) at a concentration of 0.1% by weight in RPMI 1640 (Gibco, cat. #21870-076) supplemented with 10% FBS and 1x PenStrep (Gibco, cat. #15140-122). After incubation, the samples were vortexed for 10 seconds and centrifuged at 500 g for 10 minutes at room temperature. The top layer of adipocytes and media were removed by aspiration. The remaining cells were resuspended in RBC lysis buffer, incubated for 5 minutes at room temperature and washed once in FACS buffer.

During the experiments, 25-50 µl of blood was collected from mice via tail bleed. A small incision was made on the tail, and it was gently massaged to collect the blood into tubes pre-loaded with RBC lysis buffer and kept on ice. Just before euthanasia, blood was collected via retro-orbital bleed from isoflurane-anesthetized mice. Once all samples were collected, tubes were incubated for 5 minutes at room temperature. Then, the samples were diluted with PBS containing 2% FBS and 1 mM EDTA and washed twice at 1,000 g for 10 minutes at 4°C.

### Flow cytometry

Immune cells were isolated as described above. The cells were stained with the Live/Dead Fixable Aqua Dead Cell Stain Kit (Invitrogen, cat. #L34966, diluted in PBS at 1:1000) for 5 minutes at room temperature. This was followed by staining with TruStain FcX™ PLUS (anti-mouse CD16/32) Antibody (BioLegend, cat. #156604, 1:200) and an antibody cocktail for 30 minutes at 4°C. Details of the antibodies used can be found in Table 1. Samples were washed three times in FACS buffer (PBS + 2% FBS + 1 mM EDTA). If stained for transcription factors (e.g., TOX and TCF1), True-Nuclear™ Transcription Factor Buffer Set (BioLegend, cat. # 424401) was used. In brief, after incubating with surface antibodies and three washes in FACS buffer, the cells were fixed in 100 µl fixation solution for 30 minutes at room temperature. Next, 100 µl of permeabilization buffer was added to the cells, and they were incubated for another 10 minutes at room temperature, followed by three washes in permeabilization buffer. Then, the cells were incubated with intracellular antibodies overnight at 4°C. Next day, the cells were washed three times in permeabilization buffer again. The data was acquired using BD X20, BD Symphony A3, or BD Canto II. Data was analyzed using FlowJo v10.10.0.

**Table 1.** Antibody list.

| Marker | Conjugate | Clone | Manufacturer | Cat. # |
| --- | --- | --- | --- | --- |
| Cd8a | BUV805 | 53-6.7 | BD | 612898 |
| Cd8a | BV421 | 53-6.7 | BioLegend | 100738 |
| Cd3e | BUV661 | 145-2C11 | BD | 750638 |
| Cd3e | AF488 | 145-2C11 | BioLegend | 100321 |
| Pd1 | BV711 | 29.F.1A12 | BioLegend | 135231 |
| Pd1 | BV605 | 29.F.1A12 | BioLegend | 135220 |
| Pd1 | PE-Cy7 | 29.F.1A12 | BioLegend | 135216 |
| Tox | PE | TXRX10 | invitrogen | 12-6502-82 |
| Cd44 | AF700 | IM7 | BioLegend | 103026 |
| Cd45.1 | APC/Cyanine7 | A20 | BioLegend | 110716 |
| Cd45.1 | BV785 | A20 | BioLegend | 110743 |
| Cd45.2 | PE | 104 | BioLegend | 109807 |
| Cd45.2 | PE-Dazzle594 | 104 | BioLegend | 109846 |
| Cxcr5 | Biotin | SPRCL5 | invitrogen | 13-7185-82 |
| Va2 | BUV395 | B20.1 | BD | 743834 |
| Tcf1 | AF488 | S33-966 | BD | 567018 |
| Tim3 | BV421 | RMT3-23 | BioLegend | 119723 |
| Ly108 | BB700 | 13G3 | BD | 742272 |
| Ly108 | BUV395 | 13G3 | BD | 745730 |
| Cx3cr1 | BV650 | SA011F11 | BioLegend | 149033 |
| Tigit | BV786 | 1G9 | BD | 744215 |
| Cd226 | PE | 10E5 | BioLegend | 128806 |
| Il7ra | APC | A7R34 | invitrogen | 17-1271-82 |
| Cd62l | BV785 | MEL-14 | BioLegend | 104440 |
| Klrg1 | PE-Cy7 | 2F1/KLRG1 | BioLegend | 138416 |
| Cd45 | BUV563 | 30-F1 | BD | 612924 |
| TotalSeq™-C0073 anti-mouse/human CD44 |  | IM7 | BioLegend | 103063 |
| TotalSeq™-C0004 anti-mouse CD279 (PD-1) |  | RMP1-30 | BioLegend | 109127 |
| TotalSeq™-C0301 anti-mouse Hashtag 1 |  | M1/42; 30-F11 | BioLegend | 155861 |
| TotalSeq™-C0305 anti-mouse Hashtag 5 |  | M1/42; 30-F11 | BioLegend | 155869 |
| TotalSeq™-C0310 anti-mouse Hashtag 10 |  | M1/42; 30-F11 | BioLegend | 155879 |
| TotalSeq™-C0315 anti-mouse Hashtag 15 |  | M1/42; 30-F11 | BioLegend | 155889 |

In the OT-1 x RAG2 KO experiment (Fig. 1d), staining for SIINFEKL-specific CD8+ T cells was performed. The cells were stained with the Live/Dead Fixable Aqua Dead Cell Stain Kit (Invitrogen, cat. #L34966, diluted in PBS at 1:1000) for 5 minutes at room temperature. Then, 50 μl FACS buffer with APC-labelled SIINFEKL–H-2Kb tetramer was added, and the cells were incubated at 37 °C for 30 min. Finally, an antibody cocktail was added in 50μl volume, and the staining proceeded as described above. Tetramers were obtained from the Bursky Center Immunomonitoring Laboratory at Washington University School of Medicine in St. Louis.

In LCMV infection model (Supplementary Fig. 1d), staining for gp33-specific CD8^+^ T cells was performed by adding Tetramer-APC to the antibody cocktail (MBL, cat. #TS-M512-2, 1:100 final dilution).

To stain for CXCR5 (Supplementary Fig. 1e-f), after the incubation with antibody-cocktail and three washes, cells were additionally incubated with PE/Dazzle594 Streptavidin (Biolegend, cat. #405247, 1:100 dilution) for 30’ at 4°C.

### OT-1 cells transfer and immunization

0.5-1.5*10^6 CD8^+^ T cells from young OT-1 mice (Rag2 KO Cd45.1 background) were transferred into recipient mice as described above. Next day, recipients were immunized with 100ug SIINFEKL alone (Anaspec, cat. #AS-60193-5) or 50ug SIINFEKL and 100ug poly I:C (InVivoGen, cat. #tlrl-pic). Tissues were collected 35-55 days later.

### LCMV infection

LCMV-Armstrong (LCMV-Arm) was propagated in BHK cells and viral titers were determined by a plaque forming assay using Vero cells. 2 × 10^5^ plaque-forming units (PFU) of LCMV-Arm were inoculated into mice via intraperitoneal route. Tissues were collected on day 42 post-infection. One aged mouse that failed to recover from infection, as determined by body weight loss, was excluded from the study.

### Plasma cytokine profiling

Just before euthanasia, approximately 400 µl of blood was collected via retro-orbital bleed from isoflurane-anesthetized mice. The samples were then centrifuged at 1,000 g for 10 minutes at 4°C. The plasma supernatant was carefully collected without disturbing the cell pellet and immediately frozen at −80°C. Plasma samples were then submitted to Eve Technologies for multiplex cytokine analysis using multiplex fluorescent bead assays (2x dilution, MD32). For each profiled cytokine, outliers were removed (values outside the range of 1.5 times the interquartile range (IQR) from the first or third quartile) and samples with concentration below detection limit were imputed as 0.5*minimal observed value.

### Fecal Lipocalin-2 ELISA

1-2 pellets of fresh fecal matter were acquired from TNF^+/+^, TNF^Δ69AU/+^ and TNF^ΔARE/+^ mice and stored at −80°C prior to processing. Fecal pellets were weighed and transferred into 1.5 ml Pink RINO tubes filled with zirconium oxide beads (Next Advance, cat. #PINKR5-RNA) and extraction buffer, 1X PBS + 1X Protease Inhibitor (GoldBio, cat. #GB-331-1) + 0.1% Tween-20 (Sigma Aldrich, cat. #P1379-500ML), was added at 100 mg of fecal matter per 1 mL of buffer. Pellets were homogenized utilizing a bullet blender (Next Advance Bullet Blender Storm Pro, cat. #BT24M) at impact energy speed setting 6 for a total of 7 minutes (4-minute cycle, 30 second rest period on ice, followed by a 3-minute cycle). Following homogenization, samples were spun down at 15000 rpm for 15 minutes at 4°C and supernatant was stored at −80°C until further use. Fecal lipocalin-2 levels were quantified utilizing Lipocalin-2 ELISA Kit (R&D systems, cat #DY1857-05) and reported as ng of lipocalin/g of feces.

### Distal ileum cytokine analysis

Following euthanasia with 5% CO2, mice were opened and perfused with ice cold 1X PBS. ∼0.4 cm2 of distal ileum was isolated and snap frozen in liquid nitrogen and stored at −80°C until further use. Peyer’s patches were removed from intestinal segments prior to isolation. For processing, tissues were kept on dry ice and weighed to prepare lysis buffer (MPER + 150mM NaCl and 1X Protease Inhibitor) at ∼50 µl per 10 mg tissue (Thermo Fisher Scientific, cat. #78501; Sigma-Aldrich cat. #746398-500G). Tissues were mechanically homogenized in lysis buffer with a pestle and kept on wet ice for 30 minutes prior to sonication (Qsonica Q125 Sonicator). Tissues were sonicated thrice in 3 second bursts with 40% amplitude while on ice. Homogenate was then centrifuged at 13-15,000 g for 20 minutes at 4°C. Supernatants were isolated and stored at −80°C until cytokine analysis. Cytokine levels were quantified utilizing Milliplex Mouse Th17 Premixed 25 Plex Magnetic Bead Panel (cat. #MT17MAG47K-PX25) and samples were normalized to total protein concentration of 5mg/mL based on BCA assay (Bio-Rad cat. #500-0116).

### Histology

Mice were euthanized utilizing a 5% CO2 chamber and subsequently perfused with ice-cold 1X PBS prior to tissue harvest. Intestinal tissues were separated from mesentery and flushed 2-3 times with ice cold 1X PBS followed by 4% PFA. The distal 5 cm of terminal ileum were cut longitudinally along the mesenteric border and gently rolled into a swiss roll utilizing the wooden end of a Q-tip^41^. Rolls were transferred directly into a tissue processing cassette and fixed in 4% PFA overnight at 4°C prior to paraffin embedding and sectioning. Tissue sections were subsequently stained with Hematoxylin and Eosin for histological evaluation.

### CD8^+^ T cells sorting

Enriched CD8^+^ T cells were stained with the Live/Dead Fixable Aqua Dead Cell Stain Kit (Invitrogen, cat. #L34966, diluted in PBS at 1:1000) for 5 minutes at room temperature. This was followed by staining with TruStain FcX™ PLUS (anti-mouse CD16/32 antibody, BioLegend, cat. #156604, 1:200) and an antibody cocktail for 30 minutes at 4°C. Details of the antibodies used can be found in Table 1. Finally, cells were sorted into cold PBS containing 2% FBS using BD Aria II sorter.

### IL-7 ELISA

Gonadal adipose tissue and thymus were minced into small pieces using scissors and incubated with shaking for 60 minutes at 37°C in a digestion solution containing collagenase D (Roche, cat. #11088882001) at a concentration of 0.1% by weight in DMEM/F12 (Gibco, cat. #11320082) supplemented with 10% FBS and 1x PenStrep (Gibco, cat. #15140-122). After incubation, the samples were vortexed for 10 seconds and centrifuged at 500 g for 10 minutes at room temperature. The top layer of adipocytes was collected, and media was removed by aspiration. Thymic cells, adipocytes, and the remaining pellet (stromal vascular fraction) were resuspended in 100 μl of RIPA buffer containing protease inhibitors and phenylmethylsulfonyl fluoride (all – Santa Cruz Biotechnology, cat. #sc-24948). Samples were stored at −20°C. Total protein was measured in each sample using Pierce™ BCA Protein Assay Kits (Thermo Scientific, cat. #23225). IL-7 concentration was measured using an IL-7 Mouse ELISA Kit (Invitrogen, cat. #EMIL7).

### Adipose tissue transplant

For the transplants between WT mice, the procedure was adapted from established techniques for brown adipose tissue transplants^38^. In brief, old and young CD45.2^+^ donor mice were anesthetized with ketamine/xylazine (200/40 mg/kg, i.p.). After aseptic preparation, a ventral midline incision was made to expose the visceral adipose tissue, and the perigonadal fat was carefully collected. While still under deep anesthesia, the mice were euthanized by cervical dislocation. Isolated tissue was kept in sterile ice-cold H720BSS and transplanted into the subcutaneous space of young CD45.1^+^ recipients as quickly as possible. In recipient mice, general anesthesia was induced with ketamine/xylazine (100/20 mg/kg, i.p.), and maintained with isoflurane/oxygen (1-3%/1-3 L/min) as necessary. With aseptic preparation completed, a small incision (2-4 mm) was made on the dorsal body surface. A subcutaneous pocket was then created via blunt dissection using a blunt-ended micro spatula. The donor adipose tissue, weighing 0.3-0.4 g (in 2-3 pieces), was carefully inserted into the pocket using Dumont forceps and further positioned with a blunt-ended micro spatula. The incision was closed with 1–2 simple interrupted sutures using 5-0 nonabsorbable sutures. Postoperative analgesia was administered with ketoprofen or meloxicam (5 mg/kg/day, s.c.) as needed. Sutures were removed 7–10 days after the procedure.

The transplantation procedure in experiments with AdiponectinCre/+ RosaDTAfl/+ fatless recipients was adapted from established techniques for adipose tissue grafting^37,42^. Young and old CD45.1^+^ donor mice were humanely euthanized in a CO₂ chamber in accordance with approved institutional protocol (22-0433). A ventral midline incision was performed to expose the visceral adipose depots, and perigonadal fat pads were carefully excised. The harvested adipose tissue was immediately placed in sterile, ice-cold saline to preserve tissue viability and transplanted into recipient mice as promptly as possible. In fatless CD45.2^+^ recipient mice, anesthesia was induced via inhalation of isoflurane (1–3% in oxygen at a flow rate of 1–3 L/min). The dorsal surface of each mouse was shaved and disinfected sequentially with Betadine and 70% ethanol three times. Mice were positioned on a heating pad throughout the procedure to prevent hypothermia. A small dorsal skin incision (2–4 mm) was made, and several subcutaneous pockets were created by blunt dissection using curved hemostats. Donor adipose tissue, totaling 0.3–0.4 g (divided into 2–3 fragments), was carefully inserted into each pocket and positioned with blunt forceps to ensure optimal placement while avoiding tissue puncture. The incision was closed using 2-3 simple surgical knots with 5-0 non-absorbable nylon. Postoperative analgesia was administered via subcutaneous injection of extended-release buprenorphine (1 mg/kg). Animals were monitored daily, and sutures were removed 7–10 days after the procedure.

### Lung transplant

Orthotopic left lung transplants were performed using young or old CD45.2^+^ donors and young CD45.1^+^ recipients, as previously described^43^. Transplant recipients were sacrificed at 1 month after transplantation.

### Single-cell RNA-seq: cell preparation

Splenocytes were isolated as described above, omitting the RBC lysis step. CD8⁺ T cells were enriched using the EasySep™ Mouse CD8⁺ T Cell Isolation Kit (STEMCELL Technologies, cat. #19853) according to the manufacturer’s instructions. Enriched cells were stained with Live/Dead Fixable Aqua Dead Cell Stain (Invitrogen, cat. #L34966; 1:1000 dilution in PBS) for 5 minutes at room temperature. A surface antibody cocktail (1:300 final dilution) was prepared in 1% BSA/PBS and supplemented with TruStain FcX™ PLUS (anti-mouse CD16/32, BioLegend, cat. #156604; 5 μL per sample). 25 μL of this cocktail were added to each sample, followed by a 10-minute incubation at 4°C. Subsequently, 25 μL of a TotalSeq™-C antibody cocktail (1:50 final dilution), along with a TotalSeq™-C Hashtag antibody (0.2 μg per sample), were added and samples were incubated for 30 minutes at 4°C. Details of the antibodies used are provided in Table 1. To minimize aggregate formation, all antibody cocktails were centrifuged at 14,000 × *g* for 10 minutes at 4°C prior to use. After staining, cells were washed once in 1% BSA/PBS and live CD3e⁺CD8a⁺ T cells were sorted using the Aurora Cell Sorter (Supplementary Fig. 3a). Post-sort, cells were counted and assessed for viability. The final suspension was adjusted to 1,300 cells per μL, and equal numbers of cells from up to four samples were pooled prior to library preparation.

### Single-cell RNA-seq: library preparation and sequencing

45k cell per pooled sample were subjected to droplet-based massively parallel single-cell RNA sequencing. cDNA was prepared after the GEM generation and barcoding, followed by the GEM-RT reaction and bead cleanup steps. Purified cDNA was amplified for 11-16 cycles before being cleaned up using SPRIselect beads. Samples were then run on a Bioanalyzer to determine the cDNA concentration. V(D)J target enrichment (TCR) was performed on the full-length cDNA. Gene Expression, Enriched TCR and Feature libraries were prepared as recommended by the 10x Genomics Chromium GEM-X Single Cell 5’ Reagent Kits User Guide (v3 Chemistry Dual Index) with Feature Barcoding technology for Cell Surface Protein and Immune Receptor Mapping user guide, with appropriate modifications to the PCR cycles based on the calculated cDNA concentration. For sample preparation on the 10x Genomics platform, the Chromium GEM-X Single Cell 5’ Kit v3, 16 rxns (PN-1000699), Chromium GEM-X Single Cell 5’ Chip Kit (PN-1000698), Chromium Single Cell Mouse TCR Amplification Kits (PN-1000254), Dual Index Kit TT Set A, 96 rxns (PN-1000215), Chromium GEM-X Single Cell 5’ Feature Barcode Kit v3, 16 rxns (PN-1000703) and Dual Index Kit TN Set A, 96 rxns (PN-1000250) were used. The concentration of each library was accurately determined through qPCR utilizing the KAPA library Quantification Kit according to the manufacturer’s protocol (KAPA Biosystems/Roche) to produce cluster counts appropriate for the Illumina NovaSeq6000 instrument. Normalized libraries were sequenced on a NovaSeqX plus S4 Flow Cell using the XP workflow and a 151×10×10×151 sequencing recipe according to the manufacturer protocol. A median sequencing depth of 50,000 reads/cell was targeted for each Gene Expression library and 5000 reads/cell for each V(D)J and Feature library.

### Single-cell RNA-seq: data analysis

Raw sequencing data were processed using the Cell Ranger multi pipeline (v9.0.1). Reads were aligned to the mm10-2020-A mouse reference genome for gene expression and to vdj_GRCm38_alts_ensembl-7.0.0 for V(D)J data. Sample demultiplexing was performed using antibody-derived hashtag oligos (HTOs). Filtered feature-barcode matrices were imported into Seurat^44^ (v5.1.0) for downstream analysis.

Quality control was performed with the miQC tool^45^ (via the *SeuratWrappers* R package v0.3.5) to remove low-quality cells and those with high mitochondrial gene content. Gene expression values were normalized by total UMI counts per cell, scaled to 10,000, and log-transformed. Highly variable genes (HVGs) were identified by accounting for both mean expression and dispersion. Data were scaled while regressing out technical covariates, specifically total UMI counts (nCount_RNA) and mitochondrial read fraction. Principal component analysis (PCA) was performed on the scaled data, and the first 30 principal components were used for UMAP visualization. A shared nearest neighbor (SNN) graph was constructed using FindNeighbors(reduction = “pca”, dims = 30), followed by clustering with FindClusters(reduction = “pca”, dims = 40). Cells with nCount_RNA < 5,000 and those falling into non-T cell contamination clusters were excluded, and the analysis was re-run to generate the final object. Based on clustering at a resolution of 0.1, T_AA_ cells (Cluster 2) were subset and analyzed independently using the same pipeline. Differentially expressed genes (DEGs) between clusters were identified using the Wilcoxon rank-sum test via FindAllMarkers(logfc.threshold = 0, min.pct = 0). Pathway enrichment analysis was performed using fgsea^46^ package (v1.20.0).

For TCR analysis, the scRepertoire 2^47^ R package (v2.0.4) was used. V(D)J contigs were imported using loadContigs and aggregated to the cell level with combineTCR, excluding cells with more than two receptor chains. Clonotypes were assigned to the Seurat object using combineExpression(clonalCall = “strict”), which also computed clonal frequencies and proportions. Clonal expansion within clusters was summarized using clonalOccupy, and shared expanded clonotypes were visualized with clonalCompare.

### Public gene expression data analysis

To generate the transcriptional signature of progenitor and terminal exhausted CD8^+^ T cells^27^, we used transcriptional data from GSE149876. Differential expression analysis was conducted using lmFit() and eBayes() functions from the limma package^48^ (v3.50.1), comparing progenitor exhausted CD8^+^ T cells to terminally differentiated exhausted CD8^+^ T cells. Multiple testing adjustments were made using the Benjamini-Hochberg procedure.

Tabula Muris Senis data^33^ was accessed using the TabulaMurisSenisData package (v1.0.0). Samples with total raw counts below 20,000 were removed from the analysis. Raw expression counts were then normalized using the getVarianceStabilizedData() function from DeSeq2^49^ (v1.34.0). For a cross-sectional comparison of *Il7* expression in different tissues (**Fig. 6h**), we used tissue samples from mice 6 months old and younger and mice 18 months old and older. For correlation analysis, samples of all ages were used, and Pearson’s correlation coefficient was calculated. Multiple testing adjustments were made using the Benjamini-Hochberg procedure.

Pre-processed adipose tissue snRNA-seq data^35^ (mouse_all.rds) was downloaded from https://gitlab.com/rosen-lab/white-adipose-atlas. Only chow-fed samples of perigonadal adipose tissue were used in the analysis. Clustering and annotation information was derived from the downloaded object.

### Statistics and data visualization

Statistical analysis and data visualization were performed using R (v4.1.2). The specific statistical methods used to determine significance are indicated in the figure legends. In all boxplots, the lower and upper hinges of all boxplots represent the 25th and 75th percentiles. Horizontal bars show median value. Whiskers extend to the values that are no further than 1.5 × IQR from either the upper or the lower hinge. In all crossbars and bar plots horizontal bar shows mean. Error bars represent standard error. Schematics were created with BioRender.com

## Data availability

The single-cell RNA-seq data generated in this study will be available in the GEO database. Publicly available datasets used in this study can be accessed via GEO database (GSE149876, exhausted CD8^+^ T cell subsets), TabulaMurisSenisData package (v1.0.0), and at https://gitlab.com/rosen-lab/white-adipose-atlas (adipose tissue snRNA-seq data).

## Exact group sizes

Fig. 1: (c) n=5; (d) n=2-3; (e) n=3-5; (g-h) n=4-5, (i) n=2-5 per group. Fig. 2: (b) n=17; (c, e, f) n=43; (d) n=10-19. Fig. 3: (b, c) n=5-10; (d) n=9; (h) n=3-4 per group. Fig. 4: (b) n=3-4, (h) n=3-11; (j-k) n=4-5 per group. Fig. 5: (c, g) n=3-4; (i) n=3 per group. Fig. 6: (b) n=7, (e) n=3, (f) n=17, (h) n=16, (j) n=52-56, (k) n=1-5 per group. Fig. 7: (b, d) n=3-6; (c, e) n=4 per group. Fig. S1: (b) n=3-8, (c) n=5, (d) n=3-4, (f) n=3, (h-j) n=6 per group. Fig. S2: (b) n=4-6, (c) n=44-79, (e) n=3-9, (f) n=9, (g-i) n=5-6 per group. Fig. S3: (d) n=3-4 per group. Fig. S5: (b) n=3-4. Fig. S6: (a) n=3, (d) n=5-6 per group.

