## Supplementary Figures for "Inflammaging in aged tissues drives remodeling of the CD8^+^ T cell compartment"

**Suppl. Fig. 1. Antigen and tissue exposure are required for $T_{AA}$ cell development.**

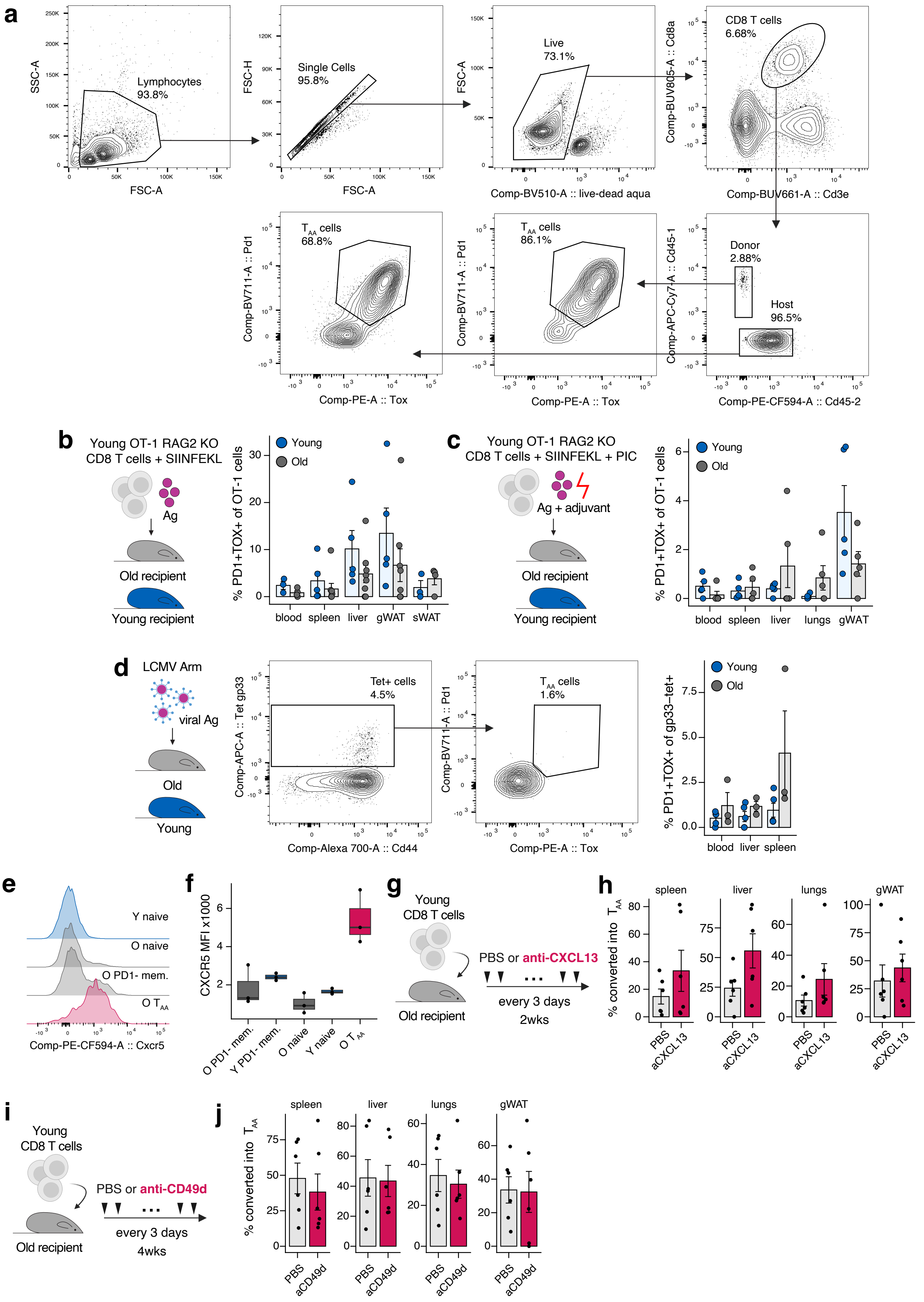

### Suppl. Fig. 1. Antigen and tissue exposure are required for $T_{AA}$ cell development.

**a.** Gating strategy used to identify *de novo*-developed $T_{AA}$ cells in the adoptive transfer model. **b-c.** Young and old mice were adoptively transferred with OT-I $CD8^+$ T cells and immunized with SIINFEKL peptide either without **(b)** or with **(c)** adjuvant. Memory OT-I cell phenotype was assessed >35 days post-immunization. **d.** Phenotype of gp33-specific $CD8^+$ T cells in young and old mice infected with acute LCMV, evaluated at day 42 post-infection. **e-f.** CXCR5 expression on corresponding $CD8^+$ T cell populations. **g.** Schematic of experimental design for CXCL13 blockade. **h.** $T_{AA}$ cell development in old mice treated with anti-CXCL13 blocking antibody or PBS control. **i.** Schematic of experimental design for CD49d blockade. **j.** $T_{AA}$ cell development in old mice treated with anti-CD49d blocking antibody or PBS control. Data are mean ± s.e.m. for **(b-d, h, j).** Exact group sizes are provided in methods.

**Suppl. Fig. 2. TNF$^{\Delta 69AU/+}$ mice serve as a model for persistent low-grade inflammation.**

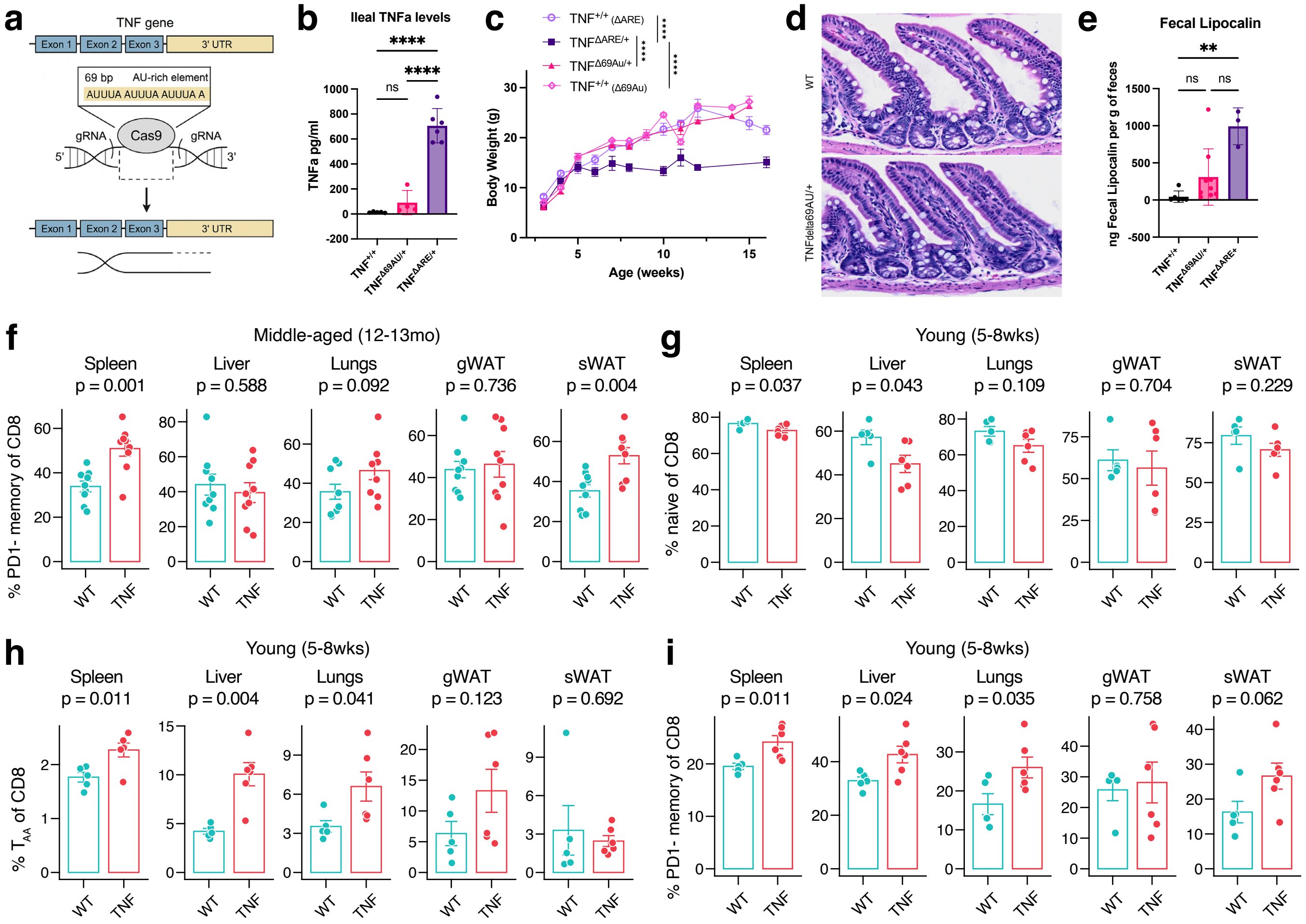


**a.** Schematic of the genetic modification at the TNF locus in TNF$^{\Delta 69AU}$ mice. For more details, see Supplementary Fig. 7. **b.** TNFa protein levels in TNF$^{+/+}$, TNF$^{\Delta 69AU/+}$, and TNF$^{\Delta ARE/+}$ mice at the terminal ileum of 13-15 weeks old females. **c.** Body weight comparison between TNF$^{+/+}$, TNF$^{\Delta 69AU/+}$, and TNF$^{\Delta ARE/+}$ mice by age (males & females combined, 3-15 weeks old). **d.** Representative images of H&E staining at the terminal ileum of 9-10 weeks old TNF$^{+/+}$ and TNF$^{\Delta 69AU/+}$ females. **e.** Quantification of Lipocalin-2 levels in stool of TNF$^{+/+}$, TNF$^{\Delta 69AU/+}$, and TNF$^{\Delta ARE/+}$ mice. **f.** Frequency of PD1$^{-}$ memory CD8$^{+}$ T cells in various tissues of middle-aged TNF$^{+/+}$ and TNF$^{\Delta 69AU/+}$ mice. **g-i.** Frequency of naïve **(g)**, $T_{AA}$ **(h)**, and PD1$^{-}$ memory **(i)** CD8$^{+}$ T cells in various tissues of young TNF$^{+/+}$ and TNF$^{\Delta 69AU/+}$ mice. Data are mean ± s.d. **(b, e)** or mean ± s.e.m. **(c, f-i).** One-way ANOVA with Tukey's post-hoc test **(b)**. Two-way ANOVA with Tukey's post-hoc test **(c)**. Kruskal-Wallis non-parametric test with Dunn's multiple comparison correction **(d)**. Unpaired two-sample Student's t-test **(f-i)**. Exact group sizes are provided in methods. ns $P > 0.05$, ** $P \leq 0.01$, **** $P \leq 0.0001$.

**Suppl. Fig. 3. Single-cell RNA-seq profiling of age-matched TNF$^{\Delta69AU/+}$ and WT mice.**

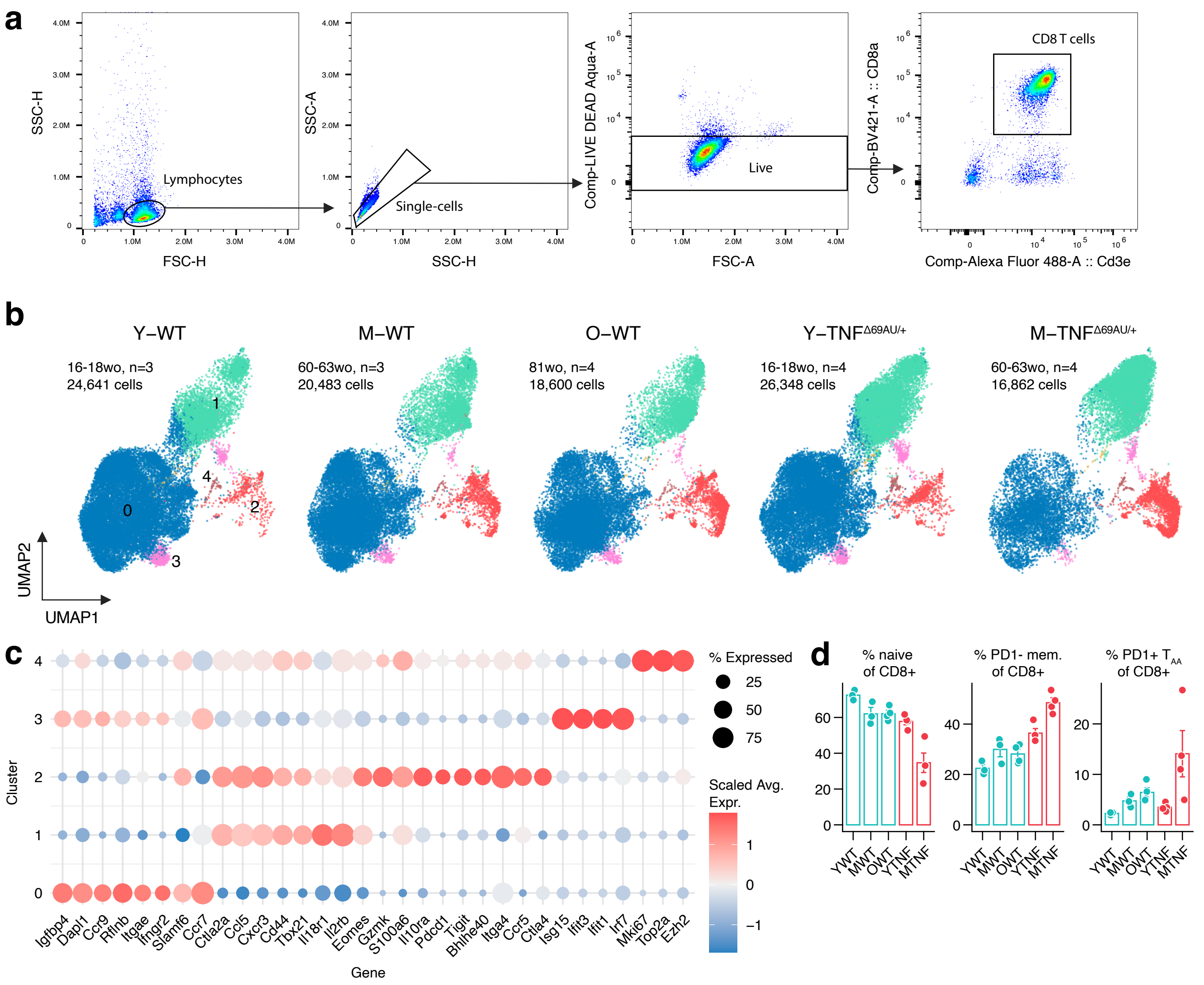


**a.** Sorting strategy used for single-cell RNA sequencing of splenic CD8$^{+}$ T cells. **b.** UMAP visualization of CD8$^{+}$ T cells isolated from WT and TNF$^{\Delta69AU/+}$ mice, split by age and genotype. **c.** Dot plot displaying scaled expression of selected marker genes across identified clusters. **d.** Flow cytometry-based quantification of CD8$^{+}$ T cell subsets in the same samples used for single-cell RNA-seq. Data in **(d)** are presented as mean ± s.e.m.

### Suppl. Fig. 4. Heterogeneity of $T_{AA}$ cells

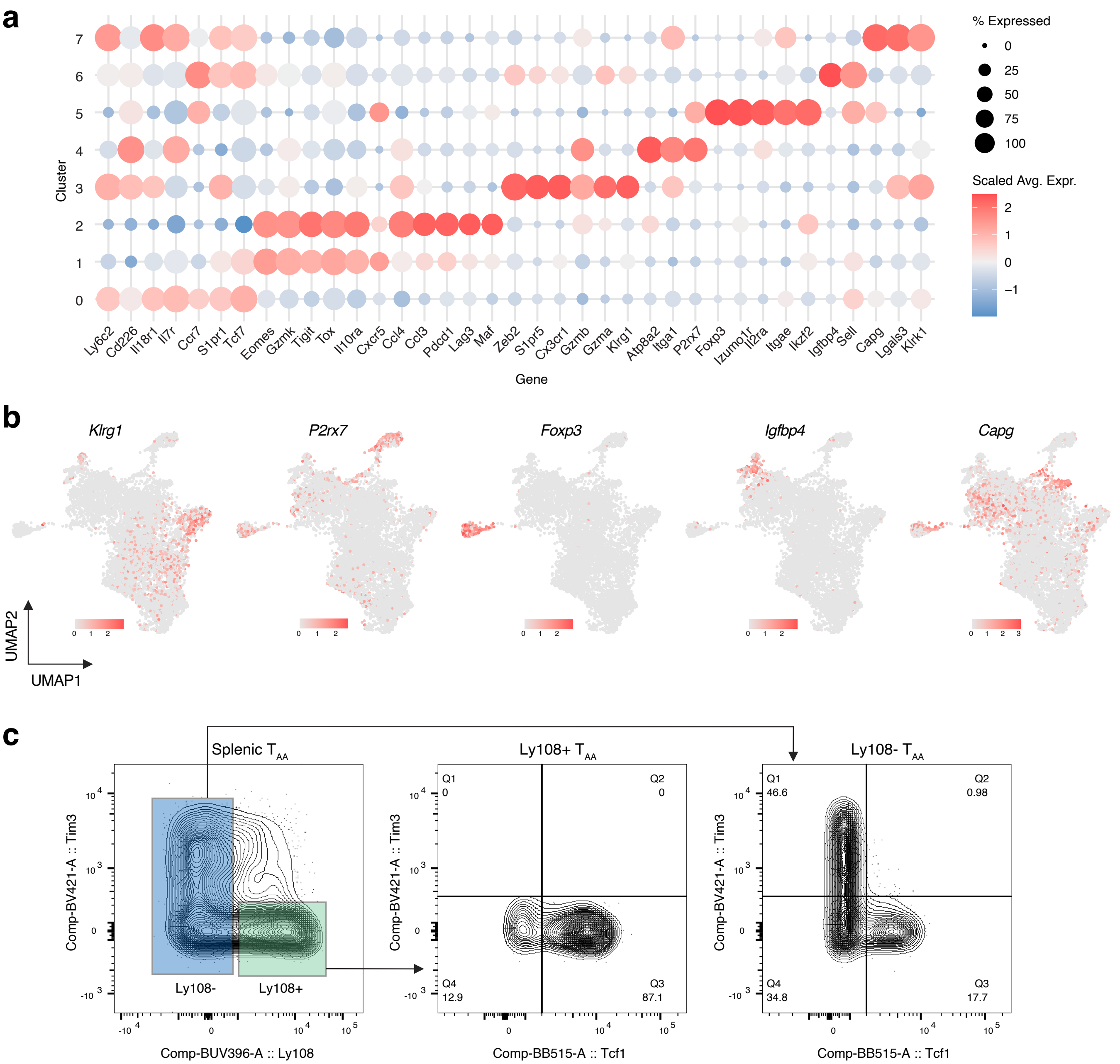


**a.** Dot plot showing scaled expression of selected marker genes across identified $T_{AA}$ subclusters. **b.** UMAP plot illustrating expression patterns of representative markers. **c.** Flow cytometry analysis of TIM3 and TCF-1 expression in $Ly108^{+}$ and $Ly108^{-}$ $T_{AA}$ cells.

### Suppl. Fig. 5. Age-associated dynamics of $T_{AA}$ cell subsets

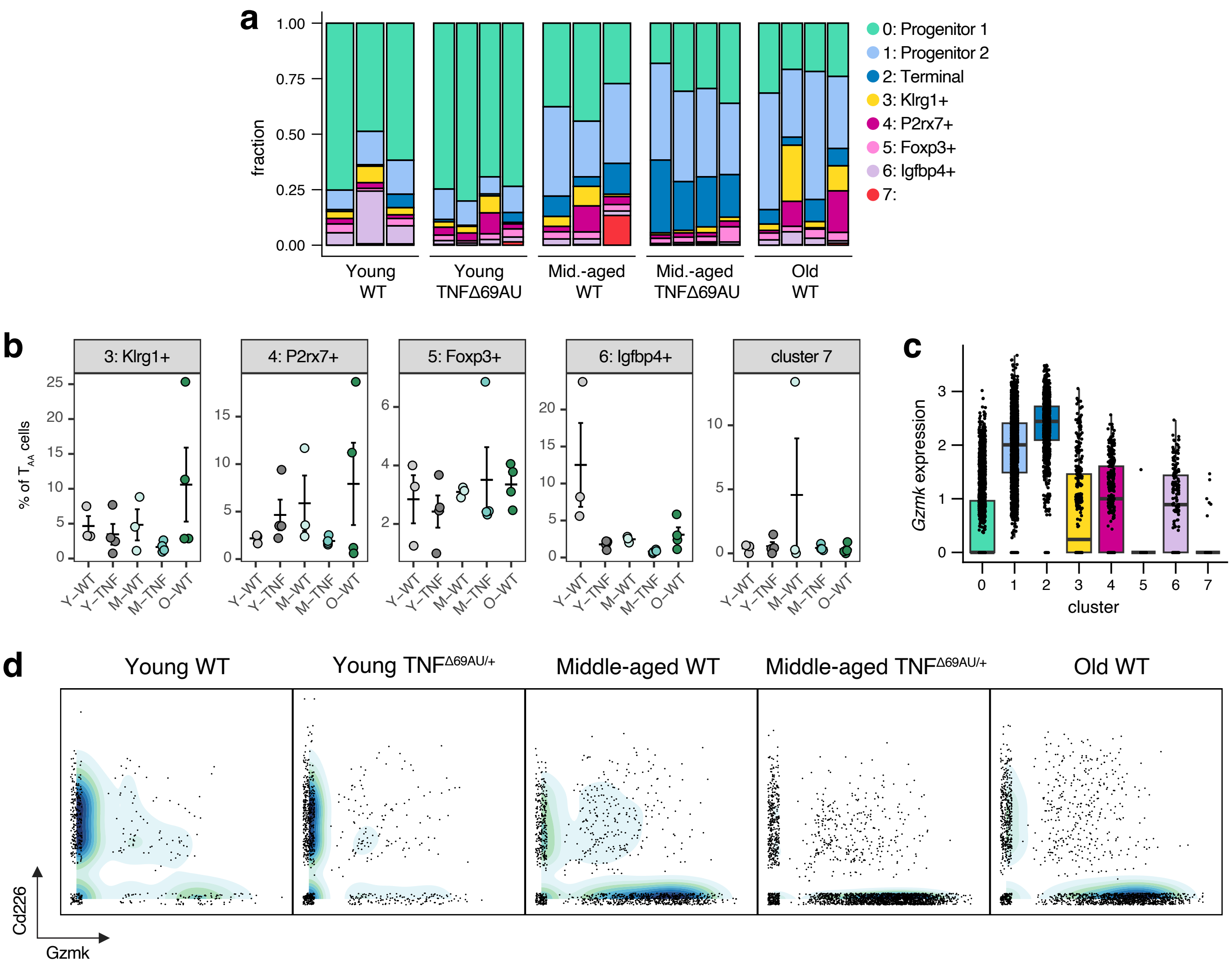


**a.** Bar plot illustrating the composition of $T_{AA}$ cells. Each bar represents an individual mouse. b. Frequency of each subcluster as a proportion of total $T_{AA}$ cells. Each dot represents an independent biological replicate. **c.** *Gzmk* expression across different $T_{AA}$ subclusters. **d.** Density plots showing the distribution of $T_{AA}$ cells based on *Gzmk* and *Cd226* expression.

### Suppl. Fig. 6. Role of adipose tissue in $T_{AA}$ cell development

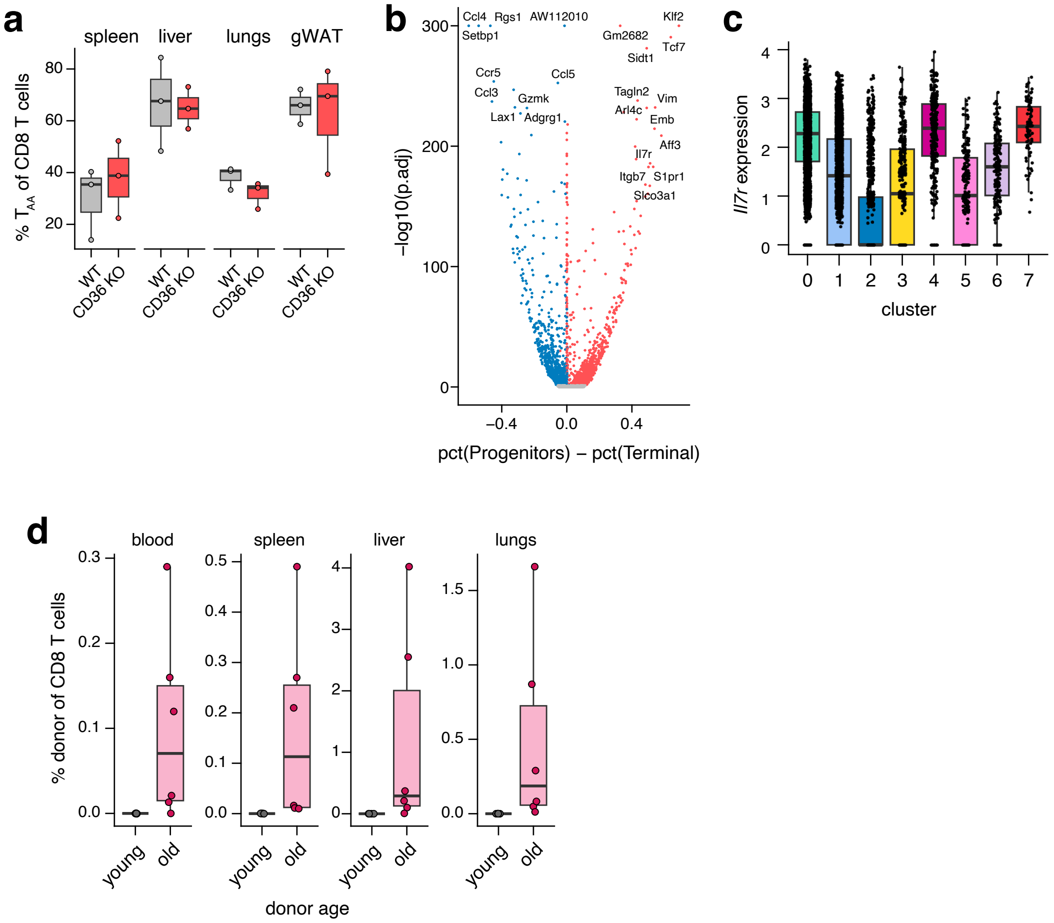


**a.** Frequency of $T_{AA}$ cells in various tissues of aged WT and CD36 KO mice. **b.** Volcano plot showing differential gene expression between Progenitor 1 & 2 and Terminal $T_{AA}$ subclusters. **c.** *Il7r* expression across $T_{AA}$ cell subclusters. **d**. Frequency of donor-derived $CD8^{+}$ T cells detected in host tissues, indicating migration from adipose transplants into peripheral organs of fatless AdiponectinCre/+ RosaDTAfl/+ recipient mice.

**Suppl. Fig. 7. Design and generation of TNF$^{\Delta 69AU/+}$ mutant mouse .**

**a**

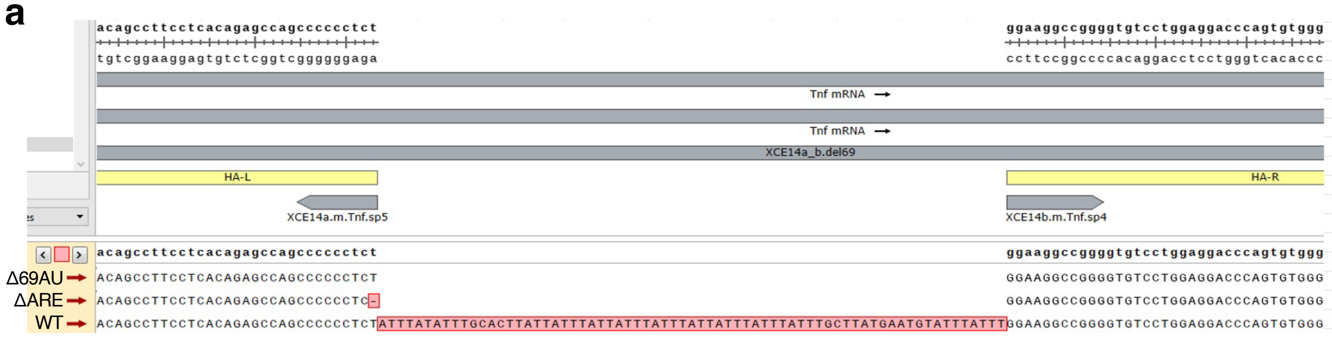


**b**

| Name | Score | Doench16 | gRNA |
|---|---|---|---|
| XCE14a.m.Tnf.sp5 | 91 | 0.698 | GTGCAAATATAAATAGAGGGNGG |
| XCE14a.m.Tnf.sp3 | 82 | 0.655 | AAGTGCAAATATAAATAGAGNGG |
| XCE14b.m.Tnf.sp4 | 88 | 0.507 | ATGTATTTATTTGGAAGGCCNGG |
| XCE14b.m.Tnf.sp5 | 88 | 0.619 | TGTATTTATTTGGAAGGCCGNGG |

**c**

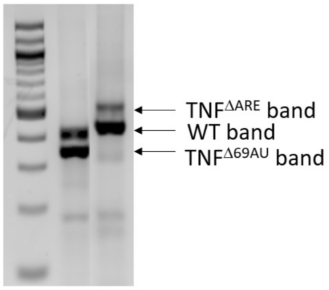


**d**

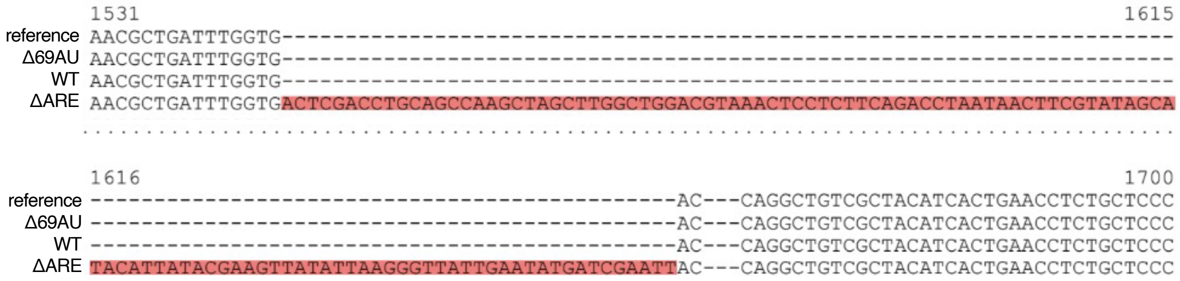


**a.** Targeting sequence and guides are schematically depicted on the upper part of the schematic. The sequencing results in the targeted locus are shown below the schematic according to genotype, with the targeted 69 bp region showing as a blank space and red highlights indicating variance of the TNFΔ69AU sequence from TNFΔARE mouse sequence or WT littermate sequence. **b.** Guide name, sequence and score for avoiding off-target effects, including the Doench 2016 score are shown. **c.** PCR products obtained during in-lab genotyping for the 3 alleles. **d.** Red highlighted sequence depicts the additional nucleotides presented within the 3' mRNA of the TNFΔARE sequence, revealing the sequence of the 116-bp addition not observed in WT or TNFΔ69AU mutants.
